# Senolytics restore hematopoietic stem cell function in sickle cell disease

**DOI:** 10.1101/2025.02.08.636742

**Authors:** Aditya Barve, Adam Cornwell, Pramika Sriram, Alex Kopyov, Preeti Dabas, Emilia Kooienga, Zakiya Kelley, James Johnson, Jacquelyn A. Myers, Esther A. Obeng, Guolian Kang, Yunus Olufadi, Terri Cain, Lindsay Talbot, David Spence, Mauricio Cortes, Sam Miller, Dirk Loeffler, Akshay Sharma, Shannon McKinney-Freeman

**Affiliations:** Department of Hematology, St. Jude Children’s Research Hospital, Memphis, TN, USA; Department of Oncology, St. Jude Children’s Research Hospital, Memphis, TN, USA; Department of Biostatistics, St. Jude Children’s Research Hospital, Memphis, TN, USA; Department of Surgery, St. Jude Children’s Research Hospital, Memphis, TN, USA; Department of Surgery, University of Tennessee Health Sciences Center, Memphis, TN, USA; Campbell Clinic, Germantown, TN, USA; Cellarity Inc., Somerville, MA, USA; Department of Bone Marrow Transplantation and Cellular Therapy, St. Jude Children’s Research Hospital, Memphis, TN, USA; Comprehensive Cancer Center, St Jude Children’s Research Hospital, Memphis, TN, USA; Department of Pathology and Laboratory Medicine, University of Tennessee, Memphis, TN USA

## Abstract

Sickle Cell Disease (SCD) is a blood disorder affecting millions worldwide. Emerging evidence reveals that SCD pathophysiology increases risk of myeloid malignancies and hematopoietic stem cell (HSC) dysfunction, possibly due to pathological stress on bone marrow. To investigate this further, we interrogated mice and individuals with SCD and observed extended cell cycle times, oxidative stress, DNA damage, senescence, and dysregulation of molecular programs associated with these processes in bone marrow hematopoietic stem and progenitor cells (HSPCs). Human SCD HSPCs displayed poor hematopoietic potential *ex vivo*. SCD mice displayed a dramatic loss of transplantable bone marrow HSPCs, which was reversed upon treatment of SCD mice with the senolytic agent, ABT-263 (navitoclax). Thus, senolytics restore bone marrow function during SCD in mice and represent a novel strategy to improve bone marrow health in individuals with SCD and improve the safety of potentially curative gene therapies that utilize autologous HSPCs from individuals with SCD.

## Introduction

SCD is an inherited hemoglobinopathy affecting approximately 100,000 people in the United States and approximately 7.74 million people globally(*1*). SCD is caused by point mutations in the β-globin subunit of the hemoglobin tetramer. These mutations cause hemoglobin to polymerize in hypoxic conditions, resulting in stiff, sickle shaped red blood cells that lyse easily and occlude microvasculature, causing acute and chronic pain, systemic inflammation, multi-organ dysfunction and early mortality(*2*). SCD epitomizes chronic hematologic stress in the form of systemic and bone marrow niche inflammation, increased erythropoietic demand and bone marrow niche damage; all insults that damage HSPCs(*3–7*). Dysfunction in HSPCs from individuals and mice with SCD is evidenced by: 1) increased numbers of circulating HSPCs(*8–10*), 2) bone marrow niche damage in SCD mice(*6, 7*), 3) bone marrow HSPCs with perturbed immunophenotypes(*8, 11–13*), 4) difficulty mobilizing HSPCs from individuals with SCD(*14*), and 5) accelerated epigenetic aging of blood mononuclear cells in individuals with SCD(*15*). Indeed, population studies of individuals with SCD report a 2-11-fold increased risk of myeloid malignancies compared to contemporary non-SCD individuals, supporting the notion of premature hematopoietic aging and genetic insult(*16, 17*). Despite accumulating evidence that the HSPC pool is progressively compromised during SCD, a definitive study of the fitness and function of HSPCs in individuals with SCD is lacking.

Allogeneic hematopoietic cell transplantation (HCT) is a potentially curative therapy for SCD but is limited by availability of human leucocyte antigen (HLA)-matched donors and immune complications associated with alternative donors(*18*). Two different autologous genetically-modified hematopoietic stem cell (HSC) products were approved for treatment of individuals with SCD in 2023 (*19, 20*). The manufacturing of these cellular products involves harvesting mobilized HSPCs via apheresis from individuals with SCD, genetically modifying them *ex vivo* and infusing them back into the patient after myeloablative conditioning. Collecting enough high-quality, long-term repopulating HSPCs from patients is critical to the success and safety of these *ex vivo* genetically modified products and remains a major rate-limiting step in their manufacturing(*14*).

Further, concern is building that SCD may select for mutated hematopoietic clones associated with elevated risk for cardiovascular and hematologic disease (*i.e.* clonal hematopoiesis), although this is currently debated (*21–24*). A recent study of HSC clonal behaviors following gene therapy observed rare HSC clones with pre-therapy mutations in *EZH2*, *DNMT3A*, *TP53* and *PPM1D* expanding overtime post-gene therapy (*25*) Further, patients with SCD who experience graft failure after allogeneic HCT are at a 40-fold increased risk of host-derived hematopoietic malignancy(*26*). These data suggest that the pathophysiology of this disease may indeed select for mutant HSPCs. Thus, as individuals with SCD live longer due to the emergence of multiple new disease modifying treatments and as allogeneic HCT and autologous gene therapies become widely available to patients, a better understanding of how HSC health and function during SCD becomes imperative (*27*).

Here, we employed flow cytometry, quantitative functional assays, and molecular profiling to investigate HSPC phenotypes and function in mice and young individuals with SCD. Our major findings include: 1) Perturbed numbers of immunophenotypic bone marrow HSPCs, 2) significant loss of long-term repopulating HSPCs, 3) prolonged cycling kinetics, elevated reactive oxygen species (ROS), DNA-damage and molecular pathways indicative of cellular stress in bone marrow HSPCs, and 4) evidence of premature senescence in the HSC pool. Our work represents the first comprehensive and quantitative functional and phenotypic examination of HSCs in mice and individuals with SCD. Importantly, treatment of young SCD mice with the senescence-targeting BCL2/BCLXL inhibitor, ABT-263 (navitoclax), increased numbers and restored HSPC long-term repopulating ability post-transplant. Thus, a premature onset of aging phenotypes correlates with loss of function in the HSPC pool of mice and individuals with SCD. We believe that this is a potential therapeutically targetable axis to improve HSPC health in individuals with SCD and improve the safety of potentially curative allogeneic and autologous HCT therapies for SCD.

## Results

### Altered frequency, diminished quiescence, and increased cellular stress in HSCs from SCD Mice

To investigate how SCD pathophysiology affects bone marrow HSPCs over time, bone marrow was isolated from young (two-months-old) and middle-aged (six-months-old) mice with SCD and interrogated by flow cytometry for HSPC frequencies (*i.e.* long-term HSCs, LT-HSCs; short-term HSCs, ST-HSCs; multipotent progenitors 2, MPP2s; MPP3s; and MPP4s) (Fig. 1A; and fig. S1A). LT-HSC frequency was modestly elevated in young SCD mice and diminished in middle-aged SCD mice, relative to non-SCD controls (Fig. 1A). ST-HSCs were similarly reduced by middle-age (Fig. 1B). Importantly, total bone marrow cellularity and MPP frequencies were unaltered in young or middle-aged SCD mice (fig. S1, B to E). An independent transgenic SCD mouse model also displayed elevated frequencies of LT- and ST-HSCs in young mice and HSC loss by middle-age (fig. S1, F and G)(*28*). No differences in apoptosis were observed between SCD and non-SCD HSCs (Fig. 1C; and fig. S2A). However, more SCD LT-HSCs and ST-HSCs were in S-G2/M and fewer were in G0 by middle-age, relative to controls (Fig. 1, D and E; and fig. S2B). Consistently, middle-aged LT-HSCs displayed elevated EdU uptake in SCD mice, relative to non-SCD controls (fig. S3A). Increased cycling correlated with an accumulation of DNA damage and ROS (Fig. 1, F to K; and fig. S3B). These data are consistent with increased replicative and cellular stress resulting in DNA damage, as previously reported(*29*).

**Figure 1.**
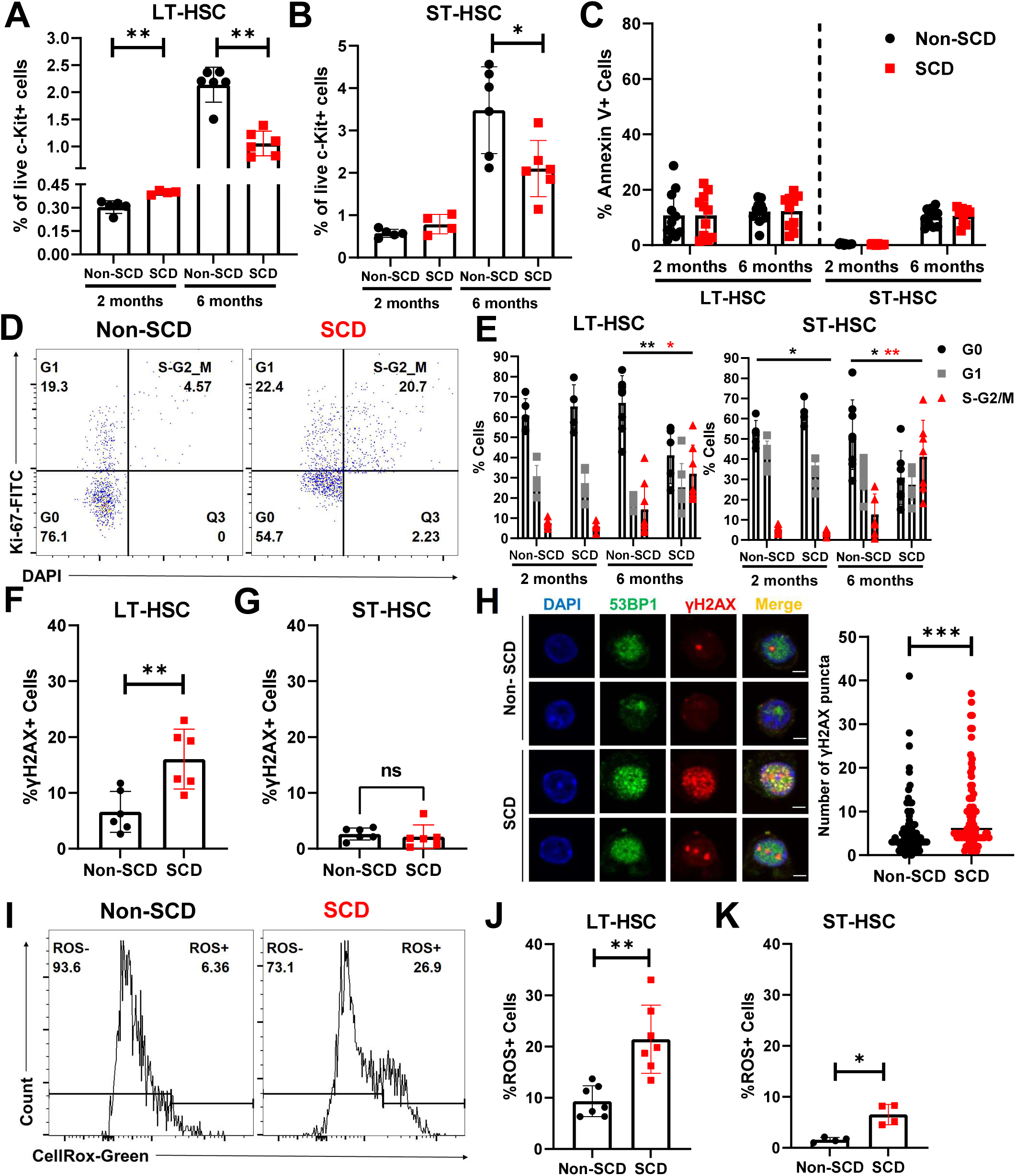
SCD perturbs HSC numbers and induces DNA damage and oxidative stress in mice LT-HSC (**A**) and ST-HSC (**B**) frequencies in young and middle-aged SCD and non-SCD littermates (n=4-6 per cohort). **C**, %AnnexinV^+^ LT-HSC and ST-HSCs in young and middle-aged SCD and non-SCD littermates (n=10-11 per cohort). **D**, Representative flow cytometry of KI-67 and DAPI in LT-HSCs of middle-aged SCD and non-SCD mice. **E,** LT-HSC and ST-HSC cell cycle status in young and middle-aged SCD and non-SCD littermates (n=5-8 per cohort). %LT-HSCs (**F**) or %ST-HSCs (**G**) with high γH2AX in middle-aged SCD and non-SCD (n=6 each) littermates. **H**, Confocal microscopy of LT-HSCs from two middle-aged SCD and non-SCD littermates (left). Co-localization of γH2AX and 53BP1 (left) and γH2AX puncta numbers/cell (right) (n=82 cells for non-SCD and n=79 cells for SCD). **I**, Representative flow cytometry of CellRox-Green in LT-HSCs of middle-aged non-SCD (left) and SCD mice (right). %ROS^+^ (**J**) LT-HSC and (**K**) ST-HSCs in middle-aged non-SCD and SCD mice (n=4-7 per cohort).

### Young and middle-aged SCD mice display loss of functional HSCs

We next quantified numbers of functional long-term blood repopulating cells (*i.e.* HSCs) in the bone marrow of young and middle-aged mice with SCD by limiting dilution transplantation(*30*). Here, 8–12-week-old lethally irradiated CD45.1^+^CD45.2^+^ C57BL/6 mice were transplanted with 5000-100,000 bone marrow cells isolated from young or middle-aged SCD mice or non-SCD controls (CD45.2^+^) along with 100,000 competitor bone marrow cells from JaxBoy mice (CD45.1^+^) (Fig. 2A). %CD45.2^+^ peripheral blood (PB) was monitored for 16 weeks post-transplant. Engraftment was defined as ≥2% total CD45.2^+^ PB and ≥1% CD45.2^+^ myeloid, B, and T cells (fig. S3c). %CD45.2^+^ PB and numbers of engrafted mice was lower in recipients of SCD bone marrow across cell doses with no evidence of lineage skewing (Fig. 2, B to E). Extreme limiting dilution analysis revealed 6.4-fold (p=0.007) and 4.2-fold (p=0.002) fewer repopulating cells in the bone marrow of young and middle-aged SCD mice, relative to non-SCD mice (Fig. 2, F and G). Thus, SCD mice display a loss of functional HSCs by two months of age.

**Figure 2.**
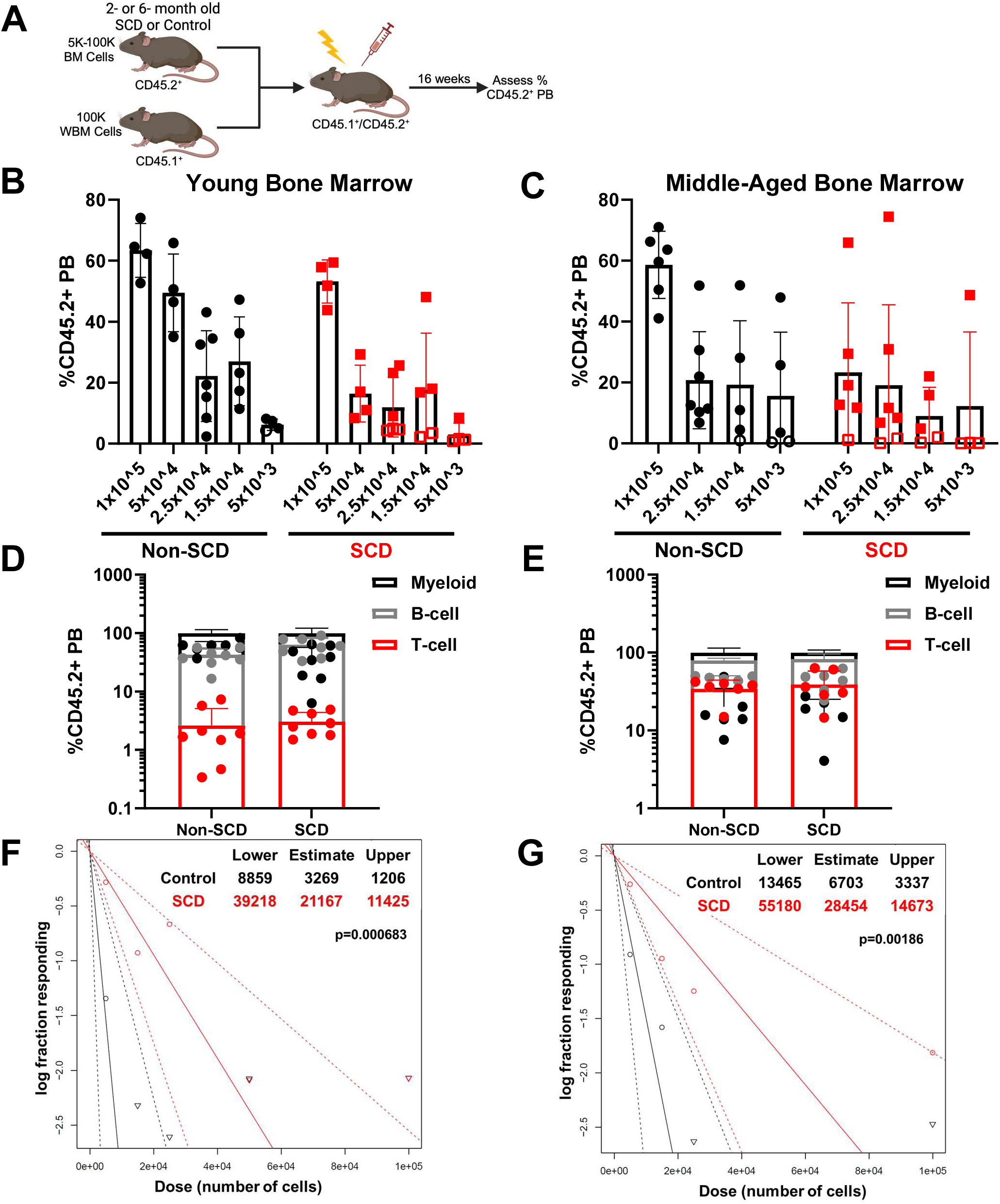
Young and middle-aged mice with SCD display a loss of functional hematopoietic stem cells. **A**) Experimental schematic of limiting dilution bone marrow transplants to estimate numbers of functional HSCs. %CD45.2^+^ PB in lethally irradiated recipients of bone marrow from young (**B**) or middle-aged (**C**) SCD or non-SCD mice (n=4-7 recipients per cohort). Cohorts receiving 5×10^3^ – 1×10^5^ bone marrow cells are shown and non-engrafted animals are represented by open symbols. %CD45.2^+^ PB in myeloid, B-cell, and T-cell lineages of engrafted recipients of 1×10^5^ and 5×10^4^ bone marrow cells from young (**D**) and middle-aged (**E**) SCD and non-SCD mice (n=4-7 recipients per cohort). Extreme limiting dilution analysis to estimate numbers of hematopoietic repopulating cells per bone marrow cell in young (**F**) and middle-aged (**G**) SCD and non-SCD mice.

### HSCs from middle-aged SCD mice display evidence of senescence-associated transcriptional and functional changes

We next performed bulk RNA-sequencing on LT-HSCs purified from the bone marrow of middle-aged SCD (n=4) and non-SCD (n=5) littermates. Principal component analysis (PCA) revealed separation between most SCD and non-SCD samples along PC1 (Fig. 3A). While correlations between replicates were >0.95, more variability was observed among SCD than non-SCD samples (fig. S4A-B). 122 and 84 significantly up or down-regulated differentially expressed genes (DEGs) were apparent in SCD versus non-SCD LT-HSCs (Fig. 3B; and fig. S4C; and Data file S1). Pathway enrichment of downregulated genes revealed downregulation of positive regulators of p53 and diminished expression of transcripts encoding ribosomal and histone proteins (Fig. 3, C and D; and fig. S4D; and fig. S5, A and B; and Data file S2).

**Figure 3.**
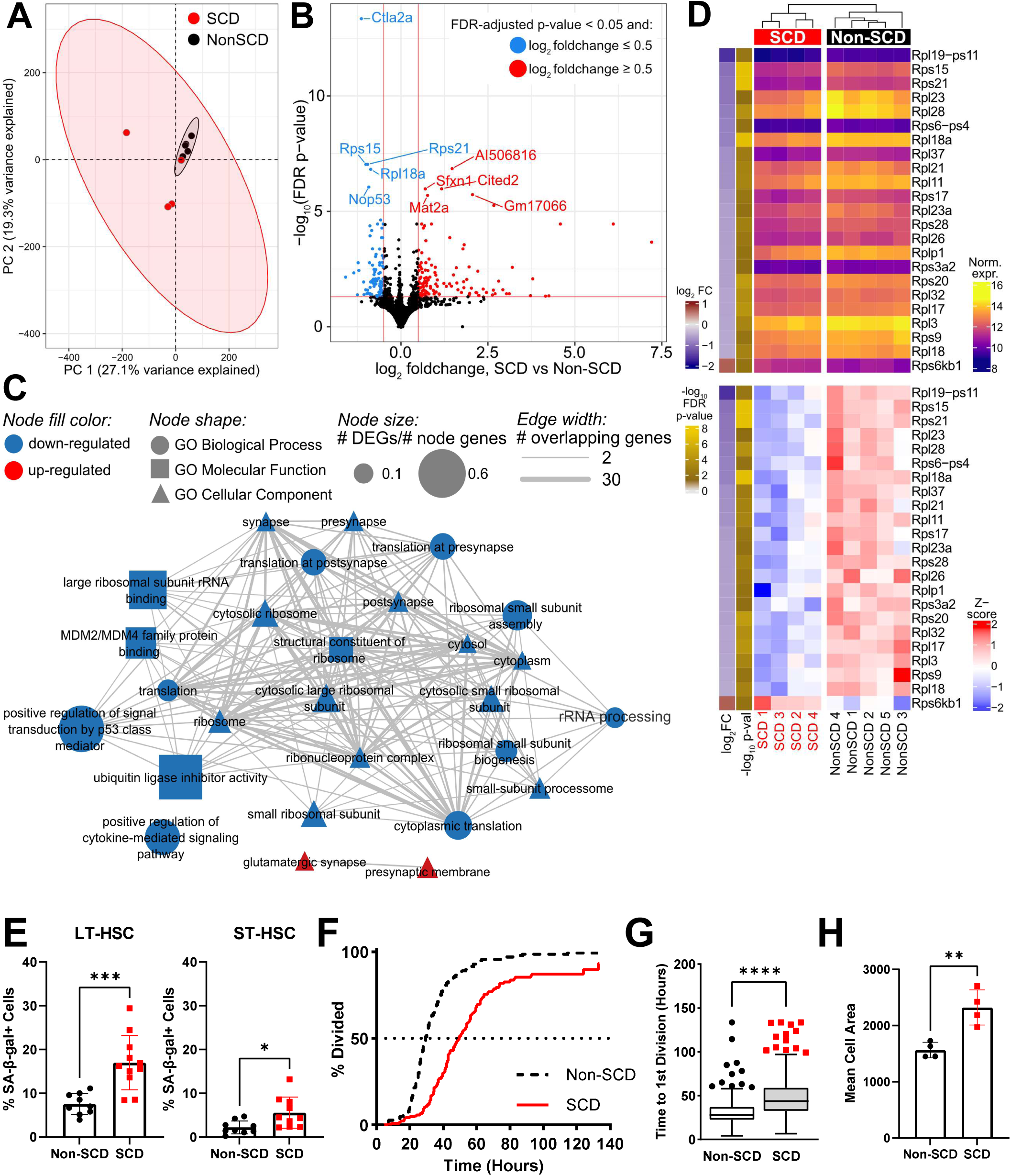
HSCs from mice with SCD display evidence of increased senescence. **A**) PCA of bulk RNA-sequencing of LT-HSCs from SCD (n = 4, red) and non-SCD (n=5, black) mice across 18,245 expressed genes. Ellipses reflect the 95% confidence interval across PC1 and PC2 for each group. **B**) Volcano plot illustrating DEGs between SCD and non-SCD LT-HSCs. Genes with FDR-corrected p-values <0.05 and log2 fold-change ≥0.5 were considered significant; horizonal and vertical red lines reflect these thresholds, respectively. Up-regulated genes denoted as red, and down-regulated genes as blue. The top 10 most significant genes by adjusted p-values are labeled. **C**) Network representation of Gene Ontology (GO) terms enriched for DEGs from (B). Enrichment based on FDR-adjusted p-values <0.05 from GOSeq for up- or down-regulated genes (red and blue nodes, respectively). Edge thickness denotes genes in common between nodes, while node size denotes ratio of No. of DEGs in dataset that fall in a term to the total number of genes associated with that term. **D**) Heatmaps of ribosome-associated protein genes with significantly altered expression in SCD mouse LT-HSCs. Left: VST-normalized expression, right: z-scores across rows to highlight the trends in expression for each gene across samples. Samples are ordered by hierarchical clustering and rows are ordered by fold-change. Adjusted p-values are also indicated in the left side bar, with brighter gold indicating lower p-values; all genes shown had FDR p-value < 0.05. **E**) Quantification of SA-β-gal^+^ LT- and ST-HSCs (left and right respectively) isolated from middle-aged SCD (n=11) and non-SCD (n=9) mice by flow cytometry. Quantification of the percent of LT-HSCs undergoing division over time (**F**), the time to first division (**G**), and mean cell area (**H**) of LT-HSCs isolated from middle-aged SCD (n=4) and non-SCD (n=4) mice at the end of the live cell imaging period.

As increased DNA damage, oxidative stress, and perturbed histone and ribosome biogenesis are often seen during senescence, we interrogated LT-HSCs for molecular and functional signatures of senescence(*31–38*). Enhanced lysosomal β-galactosidase activity outside physiologic pH (*i.e.* senescence associated β-gal, SA-β-gal) and increased cell size are conserved and well-characterized hallmarks of senescence in many cellular contexts, including HSCs(*39*). Increased SA-β-gal^+^ staining was evident in LT-HSCs and ST-HSCs from middle-aged SCD mice, relative to non-SCD mice (Fig. 3E; and fig. S5, C and D). SA-β-gal^+^ LT-HSCs were larger than SA-β-gal^-^ LT-HSCs and enriched in the SCD HSC pool, relative to controls (fig. S5E). Live imaging of cultured SCD LT-HSCs revealed marked enrichment for larger cells with slower cell cycle transition times, relative to non-SCD controls (Fig. 3, F to H; and fig. S5, F and G). Indeed, virtually no SCD HSCs underwent a division during the first 30 hours of culture, while about 50% of non-SCD HSCs divided by this time (Fig. 3F). Consistent with our analysis of fresh bone marrow, we observed comparable levels of cell death during live imaging, but significantly reduced motility, between SCD and non-SCD LT-HSCs (fig. S5, H and I). Collectively, these data suggest that SCD drives many HSCs into senescence during aging.

### Single cell transcriptomic and live imaging analysis of SCD mouse HSPCs reveals complex pool dynamics

To garner higher-resolution insight into molecular dysregulation in LT-HSCs and hematopoietic progenitors during SCD, we performed CITE-Seq on c-Kit-enriched bone marrow from young and middle-aged SCD and non-SCD mice (n=4 each). After quality filtering, 77,731 cells were annotated at the single-cell level, and validated against the CITE<u>-Seq</u> panel and marker gene expression. 22 cell types with >200 cells were identified (Fig. 4A; and fig. S6, A to C), including 765 LT-HSCs across all samples (fig. S6D). Louvain clustering segregated annotated LT-HSCs across multiple clusters, indicative of transcriptional heterogeneity (Fig. 4, B to C; and Data file S3). Ten cell clusters were unique to SCD, with two specific to young SCD mice (fig. S6E). Elevated variability amongst SCD replicates, relative to non-SCD controls, was apparent, consistent with our bulk RNA-sequencing observations (fig. S4B; and fig. S6F).

**Figure 4.**
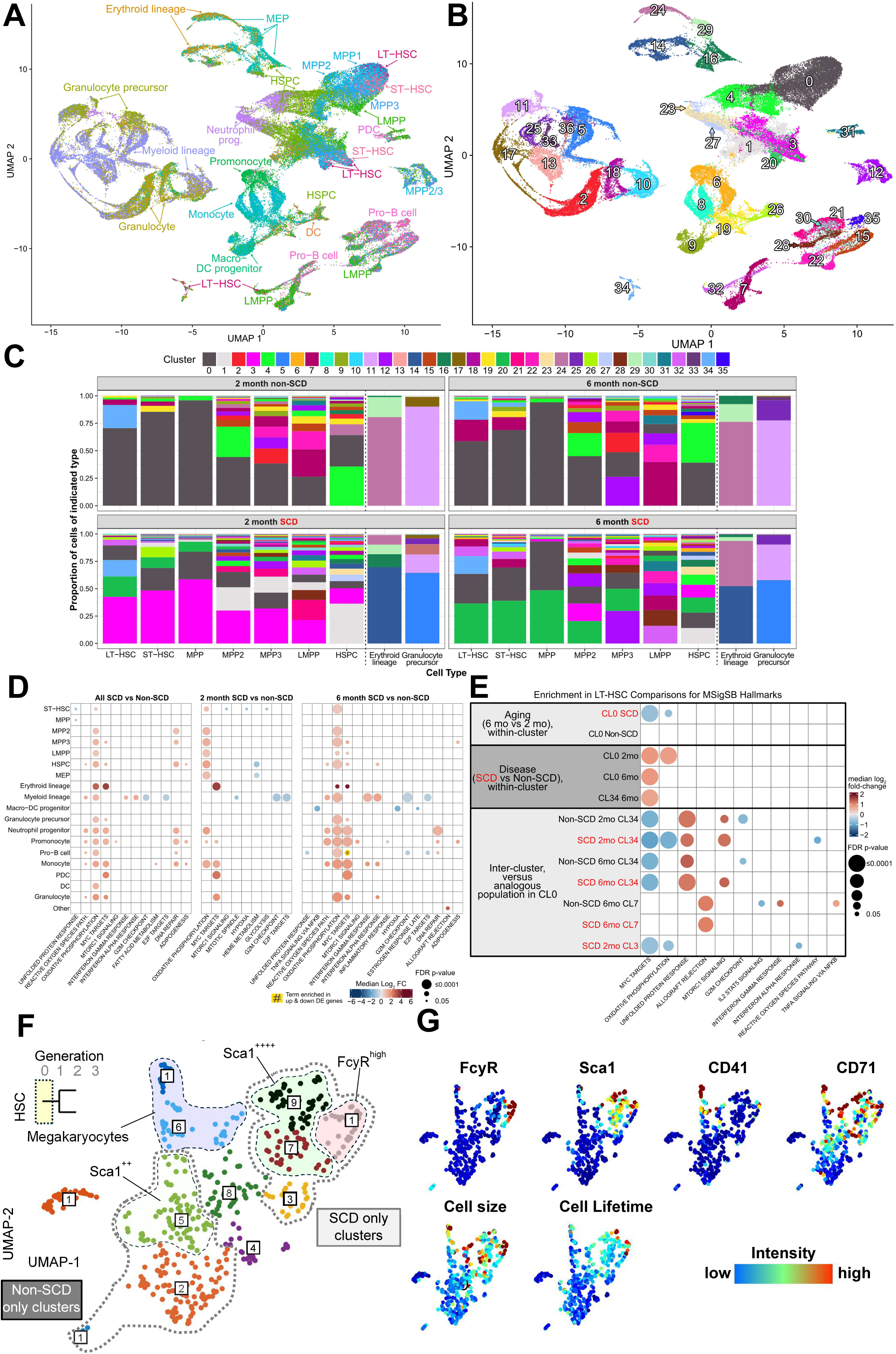
Mice with SCD display transcriptionally distinct cell populations, increased proliferation, and age-associated myeloid bias in bone marrow HSPC compartments. UMAP projections of 77,731 cells colored to highlight **(A)** annotated cell types, or **(B)** clusters from Louvain clustering. A, Cell types are indicated by label and color. B, 37 clusters were identified by unsupervised Louvain clustering at a resolution of 0.5, numbered 0-36, with CL0 containing the most cells, and CL36 the fewest. Clusters may contain multiple cell types. **C**, Distribution of cluster assignments from unsupervised clustering of early hematopoietic stem/progenitor cell types. **D**, MSigDB Hallmark gene sets significantly over-represented in pseudobulk DEGs from comparisons of all SCD vs non-SCD cells across hematopoietic progenitors (left), or young and middle-aged samples (middle and right, respectively). Circle size indicates the enrichment p-value, and color indicates the median log_2_ fold-change for the genes in the intersection with the term. Up- and down-regulated genes were run separately; in one case both were significant, as indicated by the yellow circle. Only FDR-adjusted GOSeq p-value < 0.05 results are shown. **E**, MSigDB Hallmark gene sets over-represented in DEGs from LT-HSC comparisons with MAST. Circle size indicates enrichment p-values; color indicates median log_2_ FC for the overlapping genes. The categories on the left provide the context for each of the comparison rows. **F**, Clustering of LT-HSCs by surface protein levels at the end of the live cell imaging period shows clusters unique to SCD comprised of larger, long-lived cells with high SCA-1 and FcγR (**G**).

To identify dysregulated genes due to SCD and aging amongst hematopoietic progenitors, we applied pseudobulk differential expression between SCD and non-SCD samples, in young and middle-aged animals, or with pooling across ages, for each cell type (Data file S4 to S9). DEGs upregulated by SCD were enriched for oxidative phosphorylation and MYC targets across ages throughout the hematopoietic hierarchy (Fig. 4D; and Data file S10). DEGs associated with stress responses were also apparent with aging, including an upregulated ROS response in HSPCs and myeloid progenitors, and an elevated DNA damage response; these observations are consistent with ROS-induced genomic damage during SCD. Gene expression-based cell cycle scoring revealed increased cycling of LMPPs, monocytes, and dendritic cells in young SCD mice, and a significant decline in cycling MPPs, MEPs, and erythroid progenitors with age (fig. S6G).

LT-HSCs occupied distinct age- and disease-specific clusters (Fig. 4, A and B; and fig. S6E). Their cluster distribution shifted dramatically with age and disease (Fig. 4C). We analyzed LT-HSC gene expression using MAST to highlight differences associated with disease, age, or cluster (Data file S11 to S15). Cluster 0 contained LT-HSCs from across all samples and had the largest representation in non-SCD samples and was therefore treated as a ‘baseline’ for cluster comparisons. Cluster 34 also contained LT-HSCs from both SCD and non-SCD samples. Functional enrichment for DEGs from inter-cluster comparisons revealed cluster 34 LT-HSCs as less metabolically active than those in cluster 0, regardless of disease status or age (Fig. 4E; and fig. S6H; and Data file S16 and S17). However, SCD LT-HSCs in this cluster also exhibited reduced expression of genes associated with mitochondria biosynthesis, TCA cycle, and ATP synthesis (fig. 6H), relative to non-SCD LT-HSCs. Like pseudobulk analysis, upregulated DEGs amongst SCD LT-HSCs in clusters 0 and 34 were enriched for MYC targets, relative to non-SCD LT-HSC counterparts (Figure 4E; and Data file S18). These targets included histone variant *H2az1*, cytosolic heatshock protein *Hsp90ab1*, peptidylprolyl isomerase A (*Ppia*), and the nucleophosmin, *Npm1. Ppia* promotes BCL2 activity in response to ROS, and *Npm1* regulates HSPC maintenance(*40–43*). Young SCD LT-HSCs in cluster 0 upregulated 128 unique DEGs, which suggested upregulated aerobic metabolism (*e.g.* oxidative phosphorylation and mitochondrial biosynthesis) (fig. S6, I and J). Further analyses revealed consistent upregulation of genes associated with ribosomes and translation in hematopoietic progenitors and LT-HSCs during SCD, driven by broadly higher expression of ribosome protein genes across multiple cell types and clusters (fig. S6K). Cluster seven was primarily age-associated, containing both SCD and non-SCD LT-HSCs. Upregulated DEGs associated with this cluster were enriched for immune signaling and antigen presentation (fig. S6H).

Few genes changed with age in non-SCD LT-HSCs in cluster 0. In contrast, 116 DEGs were detected when SCD LT-HSCs in this cluster were compared across ages (fig S6, L and M; and Data file S19 and S20). Most of these genes were downregulated with age yet overlapped with those upregulated overall during SCD (fig. S6N). DEGs uniquely down-regulated with age by SCD included CD74, which promotes HSC activation, sumoylation-related genes (*Serbp1*, *Sumo3*), cell cycle regulators (*Cdk4* and *Ran*), translation and RNA stability factors (*Ybx1*, *Eif4a*, *Pdcd4*, *Hnrnpd*), and epigenetic modulators (*Set*, *Ncl*, *Nap1l1*), some of which also participate in DNA repair and apoptosis (fig. S6N)(*44, 45*). These data suggest resistance to activation and a diminished DNA damage response, consistent with observed DNA damage and prolonged time to first division amongst middle-aged SCD HSCs (Fig. 1, F to H; and Fig. 3, F and G). Consistently, clustering of LT-HSCs by surface protein expression during live imaging also revealed heterogeneity amongst middle-aged SCD LT-HSCs, with unique cell clusters defined by reduced cycling, high levels of SCA-1 and FcγR, and myeloid differentiation (Fig. 4, F and G).

In sum, these data suggest that SCD promotes increased heterogeneity in the HSC pool in both young animals and during aging, the latter of which is accompanied by a global loss of metabolic activity, resistance to activation, and accelerated acquisition of aging signatures.

### HSPCs isolated from young individuals with SCD display loss of function and increased senescence

Given the observed loss of function and enrichment for senescence amongst HSCs of middle-aged SCD mice, we next interrogated bone marrow HSCs from individuals with SCD for these phenotypes. Bone marrow samples were obtained from children and young adults with SCD (n=15, ages 6 to 23 years) and without SCD (n=11, ages 2 to 21 years). HSC frequencies (Lineage^-^ CD34^+^CD38^-^CD90^+^CD45RA^-^) were elevated in individuals with SCD relative to controls, while MPP (Lineage^-^CD34^+^CD38^-^CD90^-^CD45RA^-^) and multi-lymphoid progenitor (MLP, Lineage^-^ CD34^+^CD38^-^CD90^-^CD45RA^+^) frequencies were unchanged (Fig. 5, A to C; and fig. S7A). We next examined bone marrow HSPCs (Lineage^-^CD34^+^CD38^-^) for signatures of cellular stress and senescence and elevated expression of canonical senescence mediators(*46*). DNA damage, SA-β-gal activity, and the cell cycle inhibitors, P16 and P21, were significantly increased in individuals with SCD relative to controls (Fig. 5, D to G; and fig. 7B). Thus, SCD drives elevated cellular stress and senescence hallmarks in bone marrow HSPCs of individuals with SCD.

**Figure 5.**
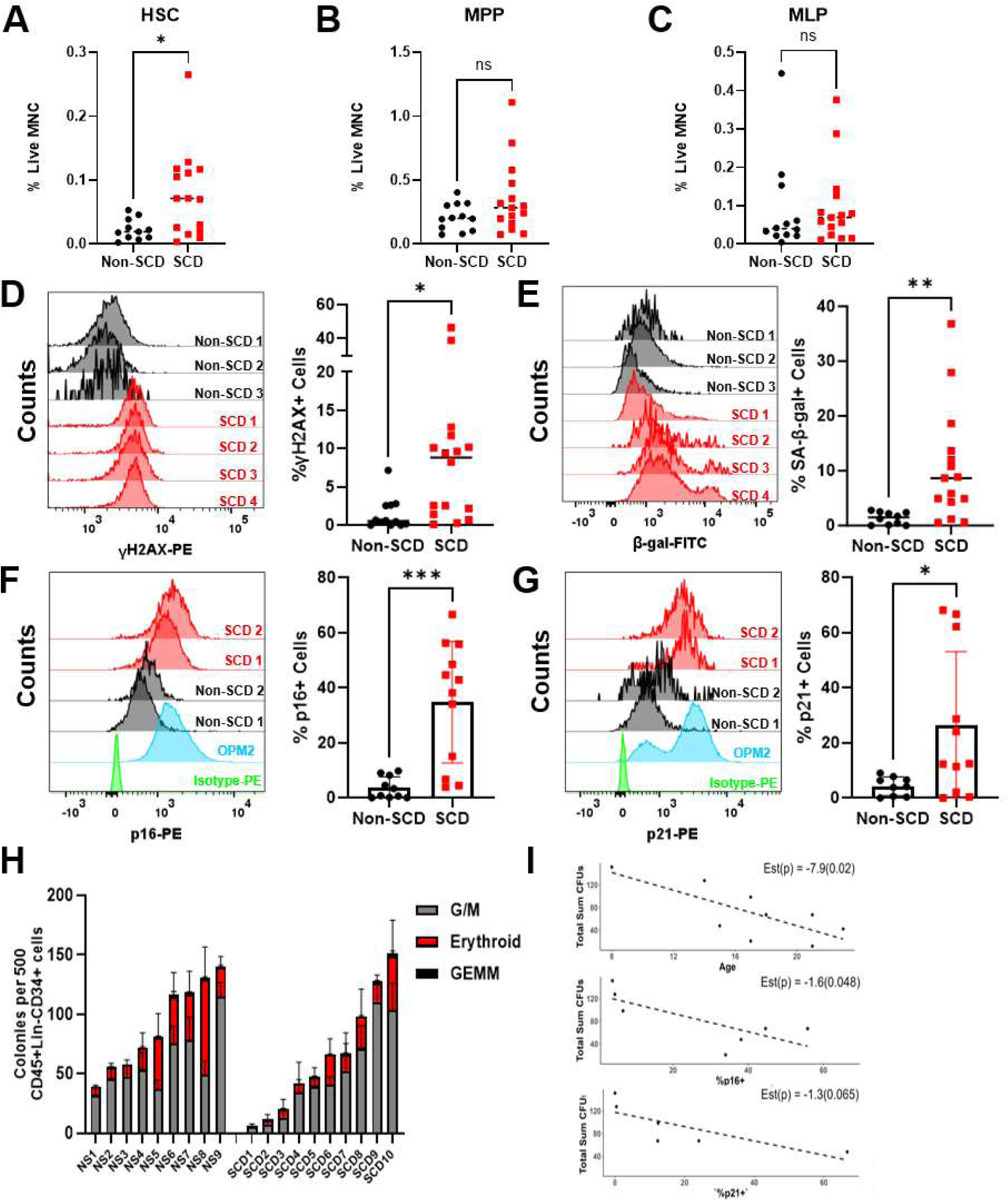
HSPC from individuals with SCD display evidence of cellular stress and loss of function. Frequency of bone marrow Lineage^-^CD34^+^CD38^-^CD90^+^CD45RA^-^ cells (*i.e.* HSCs) **(A)**, Lineage^-^ CD34^+^CD38^-^CD90^-^CD45RA^-^ cells (*i.e.* MPPs) **(B)**, and Lineage^-^CD34^+^CD38^-^CD90^-^CD45RA^+^ cells (*i.e.* MLPs) **(C)** in individuals with SCD or age-matched non-SCD controls (n=15 and 11, respectively). Representative flow cytometry plots (left) and quantification (right) of DNA damage **(D)** and SA-β-gal activity **(E)** in bone marrow Lineage^-^CD34^+^CD38^-^ cells from individuals with SCD or age-matched non-SCD controls (n=16 and 12, respectively). Representative flow cytometry plots (left) and quantification (right) of intracellular P16 **(F)** and P21 **(G)** protein levels in bone marrow Lineage^-^CD34^+^CD38^-^ cells from individuals with SCD or age-matched non-SCD controls (n=9 and 10, respectively). Isotype controls are show in green. OPM2 cells expressing P16 and irradiated OPM2 cells are shown in blue as positive controls for P16 and P21 content, respectively. **H,** CFU potential of bone marrow CD45^+^Lineage^-^CD34^+^ cells from individuals with SCD (n=10) or age-matched non-SCD (n=9) individuals. G/M: granulocyte/macrophage, GEMM: granulocyte, erythrocyte, monocyte, megakaryocyte. **I,** a significant inverse correlation between the total CFU output of HSPCs and their P16 or P21 levels was detected in individuals with SCD using a generalized linear mixed model (GLMM) with quasi-Poisson modeling.

We next interrogated the colony forming unit (CFU) potential of Lineage^-^CD45^+^CD34^+^ bone marrow HSPCs isolated from SCD and non-SCD individuals (fig. S8A; and Table S1)(*47*). SCD HSPCs displayed highly variable CFU output, relative to non-SCD controls, with some displaying very little CFU activity (Fig. 5H; and fig. S8B). Importantly, total CFU activity was inversely correlated with age, and frequencies of P16^+^ and P21^+^ bone marrow HSPCs (Fig. 5I). Myeloid potential (*i.e.* granulocyte-macrophage CFU output) also inversely correlated with %P16^+^ or %P21^+^ HSPCs (fig. S8C). In total, these data suggest that loss of function during SCD amongst bone marrow HSPCs correlates with molecular indicators of senescence.

### HSCs from individuals with SCD display transcriptional signatures of susceptibility to senescence

To better understand HSPC dysfunction in individuals with SCD, we performed bulk RNA-sequencing on Lineage^-^CD34^+^CD38^-^ bone marrow HSPCs isolated from young individuals with SCD and age-matched controls (n=7 and n=4, respectively). Each sequenced sample had at least 12,500 reliably expressed protein coding genes (counts≥10). Similarly to our observations in SCD mice, we observed substantial heterogeneity in gene expression in SCD HSPC samples relative to controls (Fig. 6A; and fig. S9A). Consequently, few DEGs were detected when all SCD samples were compared with controls (45 upregulated and five downregulated in SCD) (fig. S9B; and Data file S21). Amongst these, we observed upregulation of IL4R and IL18BP, suggesting a compensatory response to chronic inflammation, as well as TREML4, a mediator of pro-inflammatory signaling downstream of Toll-like receptors(*48–50*).

**Figure 6.**
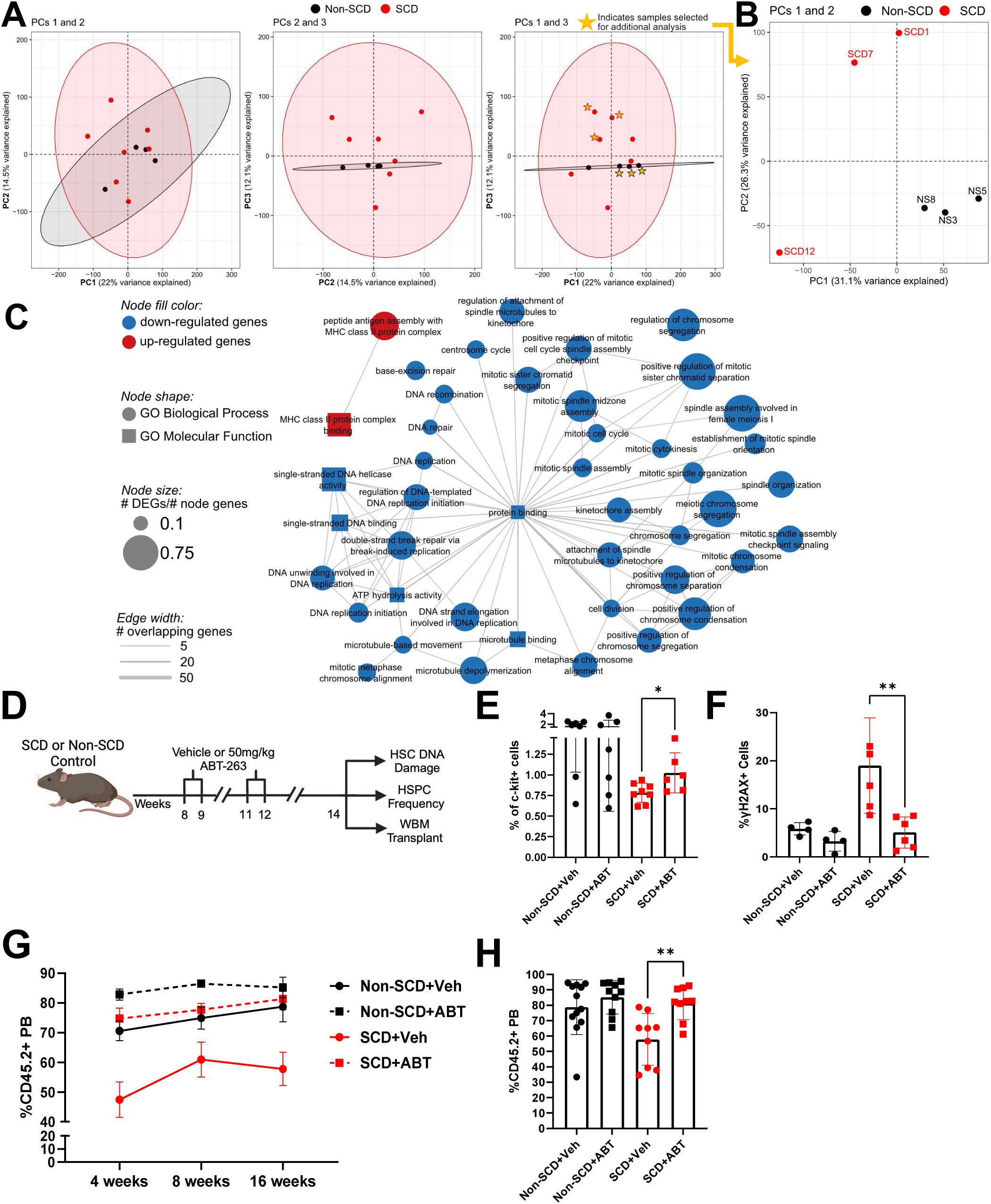
HSPCs from SCD individuals display transcriptional programs invoked by senescence and senolytic therapy restores HSPC numbers and function to mice with SCD. **A**, PCA plots for PCs 1 and 2 (left), PCs 2 and 3 (middle), and PCs 1 and 3 (right), for all non-SCD (n = 4) and SCD (n = 7) individuals across 15,699 expressed coding genes. Ellipses reflect the 95% confidence interval for each group. A subset of samples (n=3 for non-SCD and SCD) separated by condition on PCs 1 and 3, as denoted by stars. **B**, PC1 versus PC2 for the starred samples in (a). **C**, Network representation of GO terms enriched for DEGs from (B). GOSeq FDR-adjusted p-values <0.05 for up- or down-regulated genes (red and blue nodes, respectively). Edge thickness denotes the number of genes shared between terms/nodes; node size denotes ratio of the number of DEGs in the dataset intersecting a term to the total number of genes associated with a term. **D**, Treatment regimen of young SCD or non-SCD mice with ABT-263 (navitoclax) followed by bone marrow analysis and transplant to assess HSPC numbers and function. Quantification of LT-HSC frequency (**E**) and HSPC (Lineage^-^SCA-1^+^c-Kit^+^) DNA damage (**F**) in young SCD and non-SCD mice following ABT-263 treatment. **G**, %CD45.2^+^ PB of recipients of bone marrow from non-SCD or SCD mice treated with vehicle or ABT-263 overtime post-transplant (n=7-12 for non-SCD groups and n=5-9 for SCD groups). **H**, %CD45.2^+^ PB of recipients of bone marrow from non-SCD and SCD mice treated with vehicle or ABT-263 at 16-weeks post transplantation (n=10-12 for non-SCD groups and n=9 for SCD groups).

To further illuminate the molecular response to SCD pathophysiology, we next selected three SCD and three non-SCD samples that separated by condition in PC1 and PC3, although variability was still noted among the selected SCD samples (Fig. 6, A and B). 402 down- and 317 up-regulated genes were thus identified in SCD relative to non-SCD samples, including genes associated with inflammation, innate immunity, erythroid differentiation, and autophagy (fig. S9, C and D; and Data file S21). Gene network analysis revealed markedly reduced expression of genes controlling cycling, DNA synthesis, and DNA repair, highly consistent with the downstream transcriptional effects of canonical senescence induction (Fig. 6C; and Data file S22). Indeed, inhibition of TGM2 and AURKA have been linked to senescence (*51*). Genes downregulated in SCD HSPCs were significantly enriched for inhibitors of senescence in the CellAge database, with 46 overlapping genes (fig. S9E; and Data file S22). GO enrichment revealed broad down regulation of genes related to cell cycle regulation and DNA repair, whereas MHC class II genes (HLA-DRA, HLA-DOA, HLA-DMB, HLA-DBPA1, HLA-DPB1) were upregulated (fig. S9F). Thus, SCD may promote transcriptional reprogramming in HSPCs leading to a senescence-like state in some individuals.

### Senolytic treatment of SCD mice restores HSC function to bone marrow

Clearance of senescent mouse HSPCs using the BCL2/BCLXL inhibitor, ABT-263 (navitoclax), can reverse transplantation defects associated with exposure to senescence-inducing stresses, such as aging and total body irradiation(*52*). Given the observed increase in HSPCs displaying molecular and functional hallmarks of senescence in the bone marrow of mice and individuals with SCD, we tested if ABT-263 could restore HSC function and numbers to SCD mice. As HSC dysfunction manifested in young SCD mice (Fig. 2B; and Fig. 2F), we treated two-month-old SCD and non-SCD mice with two cycles of ABT-263 (50 mg/kg by oral gavage daily for two weeks followed by two weeks off) or vehicle control, as previously described (Fig. 6D)(*52*). Two weeks after final drug administration, mice were euthanized, and their bone marrow examined. An increase in LT-HSC frequency was observed in SCD mice treated with ABT-263 compared to vehicle-treated controls (Fig. 6E) and fewer HSC/MPPs (Lineage^-^Sca-1^+^c-kit^+^ cells) in treated SCD mice displayed high levels of DNA damage, relative to controls (Fig. 6F). Most importantly, ABT-263 treatment restored the hematopoietic repopulating activity of SCD mice to that of control animals with similar reconstitution kinetics (Fig. 6, G and H). These data strongly suggest that targeting senescent cells can restore function to the bone marrow during SCD.

## Discussion

We explored the impact of SCD on bone marrow HSPCs in preclinical models with age and patient samples. We found that while HSPC frequencies initially increased in young mice and patients, this increase was accompanied by elevated transcriptional and cellular heterogeneity, increased metabolic activity, increased cycling, and loss of function, including a severe loss of transplantable HSCs. HSPCs became further diminished over time during SCD, evidenced by elevated oxidative stress, DNA damage, resistance to activation, and senescence. Importantly, molecular indicators of senescence correlated with loss of HSPC function in individuals with SCD. Critically, the loss of transplantable bone marrow HSCs in SCD mice was rescued via treatment with navitoclax, a drug known to target senescent HSCs in aged mice(*52*).

HSC numbers were modestly elevated in young SCD mice but diminished with age. This loss was accompanied by prolonged cycling kinetics, ROS, and DNA damage. However, the bone marrow of young SCD mice still displayed a significant loss of blood repopulating activity in transplant studies. This is consistent with our findings in young individuals with SCD, whose HSPC frequencies were also slightly elevated, while also displaying diminished function. Inflammation is a defining characteristic of SCD that is also known to perturb the expression of key HSC surface markers(*8, 53–55*). Thus, classic cell surface markers may not reflect HSC numbers with high fidelity in mice and individuals with SCD. Indeed, single cell RNA-sequencing did not identify a difference in the overall frequency of transcriptionally defined HSCs between young and middle-age SCD mice, supporting this model and consistent with our observed loss of functional repopulating cells from SCD animals at both ages. This is also consistent with recent reports of discord between immunophenotype and transcriptionally defined HSCs when inflammatory pathways are perturbed(*56*). Importantly, unreliable HSPC immunophenotypes in SCD individuals may lead to significant overestimates of functional HSPC numbers during mobilization and apheresis collections for autologous gene therapy, resulting in manufacturing failure or delayed and poor engraftment after infusion. Indeed, a cellular product could not be manufactured for 5-10% participants in the pivotal trials for the two approved gene therapies for SCD, and this number may be higher in post-approval setting(*57–59*).

Although HSCs were reduced in SCD mice by middle-age, we paradoxically also observed an apparent increase in cycling HSCs. This may reflect functional heterogeneity amongst SCD HSCs. Indeed, LT-HSCs were distributed across multiple transcriptionally distinct cell clusters in SCD mice. These clusters were distinguished by programs for mitochondrial metabolism, post-transcriptional regulation, translation, and HSC-immune signaling. We also identified a sub-population of HSCs with SCD-specific dysregulation of key cell-cycle, activation, epigenetic, and gene expression regulatory factors in middle-aged mice, indicating a potential accumulation of damaged HSCs and disease progression by middle-age. Additionally, we also observed that a subset of SCD HSCs display a temporal lengthening of the cell cycle, which has been seen as cells transition to senescence and would read out as more ‘proliferating’ HSCs in our studies(*60, 61*). Cells transitioning to senescence are also prone to arrest in S-G2/M(*62*).

We observed broad, yet divergent, dysregulation of ribosome and translation machinery in our mouse transcriptional datasets, which were not observed in human samples. We hypothesize that this may be due to profiling of ribo-depleted total-RNA for bulk RNA-Sequencing, and poly-adenylated mRNA for 10x single cell RNA-Sequencing. This may reflect a reduction in nascent transcription and differential polyadenylation of ribosomal transcripts in HSPCs. Discrepancies could also arise from species-based differences between mice and humans in senescence presentation and/or interrogation of distinct cell pools(*63*): sorted LT-HSCs for mouse bulk RNA-Sequencing, c-Kit^+^ progenitors for mouse single cell RNA-sequencing, and heterogenous HSPCs (*i.e.* Lineage^-^CD34^+^CD38^-^ cells) for human samples. Interestingly, we observed high transcriptional variability amongst SCD samples in all datasets, reflecting the complex interactions between SCD pathology, genetics, and environment on gene expression. Increased transcriptional heterogeneity concomitant with a loss of repopulating activity is an emerging theme in recent analyses of aging HSCs(*64*).

While we did not observe transcriptional upregulation of canonical enforcers of senescence (*e.g. p16* and *p21*), we observed elevated protein levels, suggesting that their accumulation may be regulated post-transcriptionally, as previously reported(*65–69*). P16 and P21 function by inhibiting cyclin-dependent kinases that target and phosphorylate retinoblastoma protein, RB(*70*). Hypo-phosphorylated RB leads to suppression of E2F target genes, which are master transcriptional regulators of cell cycle, DNA synthesis, and DNA repair(*71*). We observed significant downregulation of these gene networks in bone marrow samples from SCD individuals, consistent with elevated P16 and P21 protein. In contrast, the only significantly upregulated gene sets in HSPCs from individuals with SCD corresponded to MHC class II genes involved in antigen presentation. These have recently been implicated in HSC regulation and may reflect antigen presentation by senescent HSPCs to facilitate culling by the immune system or active selection for the most inflammation-resistant quiescent HSPCs, which also maintain high MHC class II expression (*72, 73*). Despite high transcriptional heterogeneity, HSPCs isolated from some SCD individuals displayed conserved molecular hallmarks of senescence and cellular stress (*e.g.* elevated ROS, DNA damage, SA-β-Gal). Thus, SCD likely induces a variety of transcriptional alterations based on individual disease presentation and severity that eventually converge on senescence and impaired function. Since HSPC transcriptional signatures are highly variable during SCD, future prognostic biomarkers of SCD-induced HSPC senescence will likely require molecular, biochemical, or proteomic readouts, as opposed to transcriptional profiling.

Loss of functional HSCs from the bone marrow of SCD mice was reversed by treatment with the senolytic and BCL2/BCLXL inhibitor, ABT-263 (navitoclax). ABT-263 increased LT-HSC numbers, reduced DNA damage in the HSPC pool, and restored hematopoietic repopulating activity to SCD mice. However, the benefits of ABT-263 treatment on HSPCs likely extend beyond the originally described mechanism of action – culling of senescent HSPCs(*52*). ABT-263 can also reduce systemic inflammation by clearing cells throughout the body that may have acquired a senescence-associated secretory phenotype or via senescence-independent mechanisms(*74*). Thus, ABT-263 may improve HSPC function during SCD through both indirect and direct mechanisms, which remains to be investigated. To our knowledge, our work is the first to illustrate the therapeutic benefit of senolytics on HSPCs in SCD.

Our results are critical given the evolving landscape of potentially curative therapies for SCD. Two autologous *ex vivo* genetically modified therapies received regulatory approvals in 2023, with more in development, and advances in allogeneic HCT are making HCT safer and more accessible(*57, 58, 75, 76*). Thus, more curative treatment avenues for individuals with SCD exist now than just a few years ago, and the best and safest modality is unestablished and actively debated(*59, 77, 78*). Further, multiple new disease modifying treatments have come on the market in recent years that target different aspects of SCD pathophysiology(*27*). As these become widely adopted and available, patients are likely to live longer, paradoxically extending the period that their bone marrow is subject to the pathological stress of SCD. Understanding HSC fitness is critical to improve long-term safety and hematopoietic health following autologous genetic therapies, allogeneic HCT and throughout life.

A better understanding of HSC health during SCD might also help in patient selection and risk stratification for new therapies. Individuals with SCD are already at higher risk of developing myeloid malignancies(*16, 17*), which may partly be due to recurrent insults to HSPCs. While gene therapy might reduce or eliminate SCD pathophysiology and hence abrogate some of these insults, the genetic defects already incurred by HSCs may persist and be exacerbated during gene therapy manufacturing or hematopoietic reconstitution following infusion. HSCs harboring these genetic insults might be more susceptible to leukemic transformation due to regenerative stress, as has been observed in allogeneic HCT recipients who developed leukemia in the context of graft failure; and the observed expansion of HSC clones with driver mutations in samples from patients that underwent autologous gene therapy for SCD(*24–26*). To date, no biomarkers exist to predict outcomes and safety of genetically modified HSPCs in these patients, nor to identify individuals likely to yield an inadequate harvest of HSCs for gene therapy product manufacturing. Molecular signatures of senescence and cellular stress in HSCs may guide clinicians to not only prognosticate outcomes of prospective patients and evaluate safety of different bespoke products, but also select the best cellular therapy (allogeneic vs. autologous) for a given individual. Indeed, we observed a significant inverse correlation between loss of HSPC function and P16/P21 levels in individuals with SCD. It is noteworthy that we observed these signatures even in young individuals with SCD, thus age is not a reliable surrogate for HSC numbers or function in individuals with SCD. Further, additional benefit could come from selecting high-quality HSCs as a starting product, ideally enriching those poised to be long-lasting, durable and more genomically stable. Our proof-of-concept evaluation of senolytics to eliminate poor quality HSCs and expand functional HSCs merits further investigation in clinical studies. Together, these data provide impetus for more critical evaluation of HSC quality and function before selecting patients and cellular products for autologous gene therapy.

## Materials and Methods

### Study Design

We designed a series of studies which unveiled molecular mechanisms underlying HSPC dysfunction and demonstrated the utility of senolytics during SCD, via preclinical models and bone marrow from young individuals. Sample sizes were determined based on historical experience, no sample size calculations were performed. All replicates are included in the data presented and animals were assigned to groups based only on age and genotype (i.e. HBAA for non-SCD and HBSS for SCD) independent of sex. Animals were analyzed based on their assigned groups and covariates were controlled by using age matched non-SCD animals as controls. No specific methods were used for blinding. All statistical analysis was performed by an independent biostatistics expert who was unaware of the biological significance of distinct groups of mice or samples.

### Mice

B6;129-Hbbtm2(HBG1,HBB*)Tow/Hbbtm3(HBG1,HBB)TowHbatm1(HBA)Tow/J) (Townes model) mice were obtained from Dr. Mitchell Weiss (St. Jude Children’s Research Hospital)(*79*). C57BL/6J, C57BL/6.SJL-PtprcaPep3b/BoyJ, C57BL/6J-Ptprcem6Lutzy/J (JaxBoy) mice were obtained from Jackson Laboratory (Bar Harbor, Maine). All animals were housed in a pathogen-free facility and all experiments were carried out according to procedures approved by the St. Jude Children’s Research Hospital Institutional Animal Care and Use Committee.

### Isolation of mouse blood and bone marrow

For animal studies, bone marrow was liberated from the tibias, femurs, pelvic bones, and spines of mice by crushing followed by filtration with a 70µM cell strainer. Bone marrow cell suspensions were then incubated in red blood cell (RBC) lysis buffer (Sigma-Aldrich St. Louis, MO) for 10 minutes on ice. Mouse peripheral blood (PB) was collected from the retro-orbital sinus and RBC lysis performed as previously described(*80*). For studies assessing HSPC frequency, DNA damage by γH2AX staining, ROS burden, and SA-β-gal activity, bone marrow was isolated from mice as described but instead of RBC lysis, bone marrow was enriched for c-Kit^+^ cells (*i.e.* total HSPCs) via magnetic enrichment using anti-CD117 microbeads (Miltenyi Biotec, Carlsbad, CA) and an autoMACs magnetic cell separator (Miltenyi Biotec, Carlsbad, CA) per manufacturer’s instructions.

### Bone marrow samples and mononuclear cell (MNC) isolation from normal and SCD individuals

Bone marrow aspirates from children and young adults with SCD were acquired from the participants of an institutional bone marrow transplant protocol (NCT04362293) before undergoing conditioning and after obtaining written informed consent from the participant or their parent/guardian. All individuals with SCD who donated bone marrow were receiving hydroxyurea for a variable duration, and some received regular blood transfusions, prior to undergoing a bone marrow aspiration. This study protocol was approved by the Institutional Review Board at St. Jude Children’s Research Hospital and all study related activities were performed in accordance with the Declaration of Helsinki. Bone marrow from non-SCD individuals were acquired as clinical discard from individuals serving as healthy donors for allogeneic transplantation or undergoing orthopedic surgery for any reason.

Bone marrow MNCs were isolated by density gradient centrifugation using Ficoll-Paque PLUS (Cytiva Marlborough, MA) and centrifuging at 450 g for 30 minutes at room temperature with no brake. The MNC layer was then collected and washed twice in phosphate buffered saline (PBS) containing 2% fetal bovine serum (FBS). Total bone marrow MNCs were then resuspended in 90% FBS/10% dimethyl sulfoxide and aliquoted to a cell concentration of 1-3 x 10^6^ cells/vial and stored in liquid nitrogen.

### Preparation of mouse and human bone marrow for flow cytometry analysis

Bone marrow HSPCs were visualized by flow cytometry after staining for 20 minutes on ice with the following antibodies: Lineage cocktail [B220-BV605 (RA3-6B2), CD4-BV605 (GK1.5), CD8-BV605 (53-6.7), Gr-1-BV605 (RB6-8C5), Ter119-BV605 (TER-119)], SCA-1-PerCP-Cy5.5 (E13-161.7), c-Kit-APC-780 (2B8), CD150-PE-Cy7 (TC15-12F12.2), CD48-Alexa Fluor 700 (HM48-1) (all antibodies used at 1:200 dilution, BD Biosciences, San Jose, CA). 4’,6-diamidino-2-fenilindol (DAPI, Sigma-Aldrich, St. Louis, MO) was used for dead cell exclusion. Immunophenotypic definitions of each mouse HSPC population used throughout the manuscript are as follows: LT-HSC (Lineage^-^Sca-1^+^c-kit^+^CD150^+^CD48^-^), ST-HSC (Lineage^-^Sca-1^+^c-kit^+^CD150^-^CD48^-^), MPP2 (Lineage^-^Sca-1^+^c-kit^+^CD150^+^CD48^+^), and MPP3/MPP4 (Lineage^-^Sca-1^+^c-kit^+^CD150^-^CD48^+^).

Human cryopreserved bone marrow mononuclear cells were rapidly thawed in a 37°C water bath, washed twice and resuspended in PBS with 2% FBS and 0.1 mg/mL DNaseI. Bone marrow HSPCs were visualized by flow cytometry for HSCs, MPPs, and MLPs after staining for one hour on ice with the following antibodies: CD45-BV711 (HI30), Lineage-FITC (UCHT1; HCD14; 3G8; HIB19; 2H7; HCD56), CD34-APC-Cy7 (581), CD38-PE-Cy7 (HIT2), CD90-APC (5E10), CD45RA-PE-CF594 (HI100). All antibodies used at 1:200 dilution except CD34 antibody, which was used at 1:100 (BioLegend, San Diego CA). DAPI was used for dead cell exclusion. Immunophenotypic definitions of human HSPCs used throughout the manuscript are as follows: HSCs (Lin^-^CD45^+^CD34^+^CD38^-^CD90^+^CD45RA^-^), MPPs (Lin^-^CD45^+^CD34^+^CD38^-^CD90^-^ CD45RA^-^), and MLPs (Lin^-^CD45^+^CD34^+^CD38^-^CD90^-^CD45RA^+^).

### Flow cytometry data acquisition and analysis

For all experiments in this study, data were acquired using a 4-LASER LSR Fortessa or a 5-LASER FACSymphony™ A3 (BD Biosciences, San Jose, CA). Data analysis was performed using FloJo Version 10.8 (LLC, Ashland, OR).

### Analysis of mouse bone marrow HSPCs for apoptosis

Bone marrow cells were isolated as described above and following RBC lysis, cells were incubated on ice for 20 minutes with the following antibodies: B220-BV605 (RA3-6B2), CD4-BV605 (GK1.5), CD8-BV605 (53-6.7), Gr-1-BV605 (RB6-8C5), Ter119-BV605 (TER-119), SCA-1-PerCP-Cy5.5 (E13-161.7), c-Kit-APC-780 (2B8), CD150-PE-Cy7 (TC15-12F12.2), CD48-Alexa Fluor 700 (HM48-1). All antibodies were used at 1:200 dilutions and were from BD Biosciences (San Jose, CA). Stained cells were resuspended in Annexin V binding buffer (BD Biosciences, San Jose, CA) and then incubated with an Annexin V-FITC antibody (1:100 dilution, BioLegend, San Diego, CA) and DAPI for 20 minutes on ice before analysis by flow cytometry. Dying cells were defined as Annexin V^+^DAPI^-^.

### Cell cycle analysis of mouse bone marrow HSPCs

Bone marrow cells were isolated as described above and following RBC lysis, cells were incubated on ice for 20 minutes with the following antibodies: B220-BV605 (RA3-6B2), CD4-BV605 (GK1.5), CD8-BV605 (53-6.7), Gr-1-BV605 (RB6-8C5), Ter119-BV605 (TER-119), SCA-1-PerCP-Cy5.5 (E13-161.7), c-Kit-APC-780 (2B8), CD150-PE-Cy7 (TC15-12F12.2), CD48-Alexa Fluor 700 (HM48-1). All antibodies were used at 1:200 dilution and were from BD Biosciences (San Jose, CA). Stained cells were then fixed via incubation in PBS with 4% paraformaldehyde (PFA) for 15 minutes on ice. After washing, fixed cells were permeabilized using Perm/Wash buffer (BD Biosciences, San Jose, CA) according to the manufacturer’s instructions followed by staining with KI67-FITC (SolA15) (1:50 dilution, Invitrogen, Carlsbad, CA) and DAPI. G0 cells were defined as Ki67^-^DAPI^-^, G1 cells were defined as Ki67^+^DAPI^-^ and G2/S cells were defined as Ki67^+^DAPI^+^. Refer to Supplementary Materials for detailed information on *in vivo* 5-Ethynyl-2′-deoxyuridine (EdU) uptake study.

### Analysis of mouse and human HSPCs for oxidative stress and DNA damage

Cell ROS content was measured using the CellRox™ Green ROS detection reagent (Invitrogen, Carlsbad, CA) according to manufacturer’s instruction. Briefly, c-kit-enriched mouse bone marrow cells were incubated for 30 minutes at 37°C 5% CO_2_ with 5µM CellRox™ reagent followed by three washes and resuspension in PBS with 2% FBS. Cells were then stained for visualization of HSPCs by flow cytometry, as described above.

DNA damage was assessed in human and mouse cells by flow cytometry for phosphorylated histone γH2AX, which marks double-stranded DNA breaks (*29*). C-kit-enriched mouse bone marrow or thawed cryopreserved human bone marrow MNCs were stained for HSPC visualization, as described above. Cells were then fixed in 4% paraformaldehyde (PFA) in PBS for 10 minutes on ice followed by two washes and resuspension in 100µL of PBS with 2% FBS. Cells were then stained with an anti-γH2AX-PE (2F3, BioLegend, San Diego, CA) at 1:100 dilution for 20 minutes on ice followed by two washes and resuspension in PBS with 2%FBS. Dead cells were excluded using Live/Dead™ Aqua or Live/Dead™ Violet and cells were analyzed by flow cytometry. Refer to Supplementary Materials for detailed information on immunofluorescent microscopy DNA damage assessment studies.

### Purification of mouse and human HSPCs by cell sorting

Bone marrow was isolated from mice and enriched for c-Kit^+^ cells as described above. For collection of LT-HSCs, cells were stained with B220-BV605 (RA3-6B2), CD4-BV605 (GK1.5), CD8-BV605 (53-6.7), Gr-1-BV605 (RB6-8C5), Ter119-BV605 (TER-119), SCA-1-PerCP-Cy5.5 (E13-161.7), c-Kit-APC-780 (2B8), CD150-PE-Cy7 (TC15-12F12.2), CD48-Alexa Fluor 700 (HM48-1). DAPI (Sigma-Aldrich, St. Louis, MO) was used for dead cell exclusion.

To collect human HSPCs, cryopreserved bone marrow MNCs were rapidly thawed in a 37°C water bath, washed twice, and then resuspended in PBS with 2%FBS. For collection of HSCs, MPPs, and MLPs, cells were stained for one hour on ice with CD45-BV711 (HI30), Lineage-FITC (UCHT1; HCD14; 3G8; HIB19; 2H7; HCD56), CD34-Alexa700 (581), CD38-PE-Cy7 (HIT2). All antibodies were used at 1:200 dilution except CD34, which was used at 1:100. All antibodies came from BioLegend (San Diego CA). DAPI was used for dead cell exclusion.

For all experiments in this study, fluorescence-activated cell sorting cell sorting (FACS) was performed using a FACSAria III or FACSymphony S6 (BD Biosciences, San Jose, CA).

### Bone Marrow Transplantation

For quantitative assessment of numbers of functional HSCs, whole bone marrow (WBM) was isolated as described above from CD45.2^+^ SCD mice or non-SCD littermate controls and injected at limiting dilution (*i.e.* 5000 – 100,000 cells per recipient) into cohorts of CD45.2^+^CD45.1^+^ recipient mice via tail vein along with 100,000 CD45.1^+^ competitor WBM cells. Prior to injection, recipients were lethally irradiated with two doses of 5.5 Gy separated by 2-3 hours. Recipients were treated with 1 mL/9.1kg enrofloxacin in drinking water for 10 days post-transplant. To minimize biological variation, WBM was collected and pooled from at least three donors for all transplants. Engraftment was defined as ≥2% total CD45.2^+^ PB and ≥1% CD45.2^+^ cells in myeloid, B-cell, and T-cell compartments. Extreme limiting dilution analysis and estimation of long-term blood repopulating cells were performed using web based software as previously described, refer to Supplementary Materials for detailed information.

To assess hematopoietic repopulating potential following treatment with the senolytic, ABT-263, WBM was recovered from mice treated with drug or vehicle two weeks post final dosing. WBM marrow was pooled from three individuals from each treatment cohort and transplanted via tail vein at 2×10^5^ WBM cells/recipient into CD45.1^+^CD45.2^+^ recipients subjected to lethal irradiation along with 1×10^5^ WBM competitor cells (CD45.1^+^).

For all transplants, PB was sampled every four weeks for at least 16 weeks post-transplant and assessed for CD45.2^+^ and CD45.1^+^ PB reconstitution, as well as myeloid, B-cell, and T-cell reconstitution via staining for 20 minutes on ice with Gr1-PerCP Cy5.5 (RB6-8C5), B220-PerCP Cy5.5 (RA3-6B2), CD11b PerCP Cy5.5 (M1/70), B220-PE Cy7 (RA3-6B2), CD4-PE Cy7 (RM4-5), CD8-PE Cy7 (53-6.7), anti-CD45.2 V500 (104) and anti-CD45.1 FITC (A20). All were used at 1:200 dilution and acquired from BD Biosciences (San Jose, CA). DAPI (Sigma-Aldrich, St. Louis, MO) was used for dead cell exclusion.

### Analysis mouse and human cells for β-galactosidase activity

Senescence-associated *β-galactosidase* (SA-β-Gal) activity in mouse and human HSCs and HSPCs was assessed via a live cell flow cytometry-based assay kit (Catalog #35302; Cell Signaling Technology, Danvers, MA) according to the manufacturer’s instructions. Briefly, c-kit-enriched mouse bone marrow or thawed cryopreserved human bone marrow MNCs were incubated with 100nM bafilomycin A for one hour at 37°C and 5% CO_2_ to alkalize cellular lysosomes and inhibit autophagy. Next, cells were treated with 33µM of the provided cell permeable fluorogenic substrate (which fluoresces when hydrolyzed by β-galactosidase) for 3-4 hours followed by two washes of PBS with 2% FBS. Cells were then stained with appropriate antibodies for HSC or HSPC visualization, as described above. Dead cells were excluded using either Live/Dead™ Aqua or Live/Dead™ Violet (Invitrogen, Carlsbad, CA). Mouse LT-HSC SA-β-Gal activity was also assayed and confirmed by staining of cytospin preparations, refer to Supplementary Materials for detailed information.

### CFU potential of human HSPCs

Cryopreserved bone marrow MNCs were thawed in a 37°C water bath and transferred to a 15 mL conical tube. 5 mL of PBS with 2% FBS 1%. PEST was then added dropwise. MNCs were washed twice and resuspended in PBS with 2% FBS 1% PEST. Cells were stained with CD45-BV711 (HI30), Lineage-FITC (UCHT1; HCD14; 3G8; HIB19; 2H7; HCD56), CD34-APC-Cy7 (581), CD38-PE-Cy7 (HIT2) at 1:200 dilutions. All antibodies came from BioLegend (San Diego CA). DAPI (Sigma-Aldrich, St. Louis, MO) was used for dead cell exclusion. 500 CD45^+^Lineage^-^ CD34^+^ cells were sorted into each well of a 96-well U-bottom plate (Corning, #351172) containing 100 uL of X Vivo-10 media (Lonza, # BEBP02-055Q) with 1%BSA, 1%PSG, hGM-CSF (10 ng/uL), hTPO (15 ng/uL), hIL-6 (10 ng/uL), hFlt3L (100 ng/uL), and hSCF (100 ng/uL). All cytokines were purchased from Peprotech (Cranbury, NJ). Well contents were then transferred to 2 mL aliquots of H4435 methylcellulose media (StemCell Technologies, #04435), mixed well, and dispensed into SmartDish 6-well plates (StemCell Technologies, #27370). After 14 days incubation at 37°C with 5% CO_2_, wells were imaged with the STEMvision imager (StemCell Technologies) and colonies scored.

### Treatment of SCD mice with the senolytic, ABT-263

Cohorts of two-month-old SCD or non-SCD littermates were administered 50 mg/kg of ABT-263 (Biovalley, Nanterre, France) dissolved in 60% Phosal 50/30% PEG400/10% EtOH or vehicle by daily oral gavage for one week. Mice were then rested for two weeks, followed by another week of daily ABT-263 or vehicle via oral gavage. Two weeks later, mice were euthanized, and bone marrow collected for analysis of HSPCs and transplantation studies (detailed above).

### Advanced Analyses

For details on small input Western blot, RNA sequencing and associated data analysis, or time-lapse live cell imaging please refer to Supplementary Materials and Methods.

### Statistical Analysis

Data normality was checked using the Shapiro–Wilk test. A two-sample t-test or (exact) Wilcoxon rank sum test was used to evaluate two-group differences in clinical factors variables depending on the normality of the data (Figs 1a, 1b, 1c, 1e, 1f, 1g, 1h, 1j, 1k, 2b, 2c, 2d, 2e, 3f, 5a, 5b, 5c, 5d, 5e, 5f, 5g, 6e, 6f, 6g, and 6h). The false discovery rate correction was adjusted for multiple comparisons whenever relevant. All p values were two-sided. A linear regression model and a generalized linear mixed model (GLMM) with a quasi-Poisson model were used to compare the cases to controls with subject ID as a random effect (Fig 5I), adjusting for the number of records. Overdispersion was checked before we conducted the GLMM.

## Supporting information

All raw gene expression data from this study.

## List of Supplementary Material

### Materials and Methods

Fig. S1 to S9

Table S1

Data files S1-S22

Data file S1: Bulk RNA-Seq differential expression analysis for HSCs from middle-age SCD animals compared to middle-age non-SCD control animals.

Data file S2: Functional enrichment for DEGs from HSCs from middle-age SCD animals compared to middle-age non-SCD control animals.

Data file S3: Distribution of single-cell RNA-Seq HSPCs across clusters from unsupervised cluster analysis.

Data file S4: Mouse scRNA-Seq pseudobulk differential expression analysis for young SCD versus young non-SCD control animals, for each cell type.

Data file S5: Mouse scRNA-Seq pseudobulk differential expression analysis for middle-age SCD versus middle-age non-SCD control animals, for each cell type.

Data file S6: Mouse scRNA-Seq pseudobulk differential expression analysis for all SCD versus all non-SCD control animals, with ages pooled, for each cell type.

Data file S7: Mouse scRNA-Seq pseudobulk differential expression analysis for young versus middle-aged animals, with condition (SCD and non-SCD) pooled. for each cell type (i.e. aging irrespective of SCD status).

Data file S8: Mouse scRNA-Seq pseudobulk differential expression analysis for young SCD versus middle-age SCD animals, for each cell type (i.e. aging in SCD).

Data file S9: Mouse scRNA-Seq pseudobulk differential expression analysis for young non-SCD controls versus middle-age non-SCD control animals, for each cell type (i.e. aging in non-SCD controls).

Data file S10: Functional enrichment analysis for pseudobulk differential expression analyses from tables S4 to S9.

Data file S11: Single-cell differential expression analysis for LT-HSCs, from inter-cluster comparisons (relative to cluster 0) for non-SCD control animals.

Data file S12: Single-cell differential expression analysis for LT-HSCs, from inter-cluster comparisons (relative to cluster 0) for SCD animals.

Data file S13: Single-cell differential expression analysis for LT-HSCs, from comparisons between SCD and non-SCD controls.

Data file S14: Single-cell differential expression analysis for LT-HSCs, from comparisons between middle-age and young non-SCD control animals (i.e. aging in non-SCD LT-HSCs).

Data file S15: Single-cell differential expression analysis for LT-HSCs, from comparisons between middle-age and young SCD animals (i.e. aging in SCD LT-HSCs).

Data file S16: Functional enrichment analysis for DEGs in Table S11 (LT-HSC inter-cluster comparisons for non-SCD controls)

Data file S17: Functional enrichment analysis for DEGs in Table S12 (LT-HSC inter-cluster comparisons for SCD animals)

Data file S18: Functional enrichment analysis for DEGs in Table S13 (LT-HSC SCD vs non-SCD control comparisons)

Data file S19: Functional enrichment analysis for DEGs in Table S14 (LT-HSC aging in non-SCD controls)

Data file S20: Functional enrichment analysis for DEGs in Table S15 (LT-HSC aging in SCD animals)

Data file S21: Differential expression analysis for bulk RNA-Seq of HSCs from pediatric individuals with and without SCD

Data file S22: Functional enrichment for DEGs from human bulk RNA-Seq in Table S21. References (81-131, only cited in the SM)

## Acknowledgements

We thank the patients, parents, participating clinicians, and institutional core facilities staff (Animal Resource Center, Flow Cytometry and Cell Sorting, Experimental Hematology Flow Cytometry, and Biostatistics) at St. Jude Children’s Research Hospital without whom this work would not be possible. In particular, we thank David Cullins for his expertise in cell sorting, as well as Jenna McCommon, Amanda George, Anna Baird, Amanda Shellhart, Clemothy Bell, and Shelby Patrick for their help with mouse procedures. We also thank Charles Sherr, Mitch Weiss, and members of the Division of Experimental Hematology at St. Jude Children’s Research Hospital, for valuable input and feedback on the study and critical reading of the manuscript. This work was supported by the following grants: National Institutes of Health grant F32HL164095 (AB) National Institute of Health grant R01HL174644 (DL) National Institute of Health grant F31HL170678 (TC) National Institute of Health grant U01HL163983 (AS) National Institute of Health grant R01HL168893 (SMF) National Institute of Health grant T32CA236748 (AC) American Lebanese Syrian Associated Charities (ALSAC) (SMF, AS, DL, EAO, LT, GK) Leukemia & Lymphoma Society Scholar (SMF) Alex’s Lemonade Stand (Grant # 22-27080) (DL) Discovery Research Grant (Edward P. Evans Foundation) (EAO) American Cancer Society (RSG-22-023-01-CDP) (EAO) The content of this manuscript is solely the responsibility of the authors and does not necessarily represent the official views of the National Institutes of Health.

## Author Contributions

AB designed, performed, and analyzed most of the experiments and wrote the manuscript. AC and JAM. performed all bioinformatics analyses and AC wrote the manuscript. PS performed all human colony forming assays. PD performed immunofluorescence and frequency analyses. EK, ZK, AK, JJ, and TC also performed experiments and maintained mouse colonies. EAO, LT, and DS were involved in human sample acquisition, data interpretation, and edited the manuscript. GK and YO performed all statistical analyses. MC and SM performed all mouse single cell RNA sequencing studies. AB, DL, and AK performed and analyzed all live cell imaging studies. AS and SMF analyzed and interpreted data, wrote the manuscript, and jointly supervised the study. All authors discussed the results and provided commentary on the manuscript.

## Completing Interests

AS has received consultant fees from Spotlight Therapeutics, Medexus Inc., Vertex Pharmaceuticals, Sangamo Therapeutics, Editas Medicine, BioLineRx and Pfizer. He is a medical monitor for a Resource for Clinical Investigations in Blood and Marrow Transplantation Conditioning SCID Infants Diagnosed Early clinical trial, for which he receives financial compensation. He has also received research funding from CRISPR Therapeutics and honoraria from Vindico Medical Education and Blackwood CME. AS is the St Jude Children’s Research Hospital site principal investigator of clinical trials for genome editing of sickle cell disease sponsored by Vertex Pharmaceuticals/CRISPR Therapeutics (NCT03745287), Novartis Pharmaceuticals (NCT04443907), and Beam Therapeutics (NCT05456880). The industry sponsors provide funding for the clinical trial, which includes salary support paid to AS’s institution. AS has no direct financial interest in these therapies.

## Data and Materials Availability

The mouse bulk, single-cell RNA sequencing, and human bulk RNA sequencing datasets generated during this study have been deposited to GEO as a SuperSeries under accession number GSE277684. All other data and/or materials will be made available upon request.

## Supplementary Methods

### In vivo 5-Ethynyl-2′-deoxyuridine (EdU) uptake in mouse HSPCs

Middle-aged SCD and non-SCD mice were administered 200 mg/kg EdU by tail vein injection. 24 hours later, mice were euthanized and bone marrow was isolated and enriched for c-Kit^+^ cells, and then incubated on ice for 20 minutes with the following antibodies: B220-BV605 (RA3-6B2), CD4-BV605 (GK1.5), CD8-BV605 (53-6.7), Gr-1-BV605 (RB6-8C5), Ter119-BV605 (TER-119), Sca-1-PerCP-Cy5.5 (E13-161.7), c-Kit-APC-780 (2B8), CD150-PE-Cy7 (TC15-12F12.2), CD48-Alexa Fluor 700 (HM48-1). All antibodies were used at 1:200 dilution and were from BD Biosciences (San Jose, CA). EdU incorporation into DNA was visualized by click chemistry using the *In Vivo* EdU Flow Cytometry Kit 488 (Sigma-Aldrich, Burlington, MA) per manufacturer’s instructions and HSPC EdU uptake was analyzed by flow cytometry as described above.

### Immunofluorescence and microscopy to assess DNA damage in mouse LT-HSCs

LT-HSCs were isolated from six-month-old SCD or non-SCD littermates as described above. Coverslips were coated with 10 μg/mL CD44 in PBS for one hour at room temperature. After a gentle PBS wash, approximately 700 LT-HSCs were added to coverslips and incubated at 37°C for one hour. Cells were gently washed again and crosslinked with 4% paraformaldehyde (Electron Microscopy Sciences, Hatfield, PA) in HEPES-KOH (pH 7.5) at room temperature for 15 minutes. After three washes, cells were permeabilized and blocked simultaneously with 0.3% Triton X-100 in blocking buffer (0.5% bovine serum albumin, 50 mM ammonium chloride in PBS) for 30 minutes. Cells were then washed three times and stained sequentially with mouse anti-γH2AX (Ser139, 1:200 dilution, Sigma, Burlington, MA) and rabbit anti-53BP1 (1:200, Novus Biologicals, Centennial, CO) in blocking buffer at room temperature for one hour each. After staining, cells were washed three times and then incubated with goat anti-mouse Alexa Fluor 647 (1:300 dilution, Abcam, Waltham, MA) and donkey anti-rabbit Alexa Fluor 488 (1:300, Abcam, Waltham, MA) in blocking buffer at room temperature for one hour. Coverslips were mounted on slides using ProLong Diamond with DAPI (Thermo Fisher, Waltham, MA). Images were captured using a Leica SP8 confocal microscope at 63X objective lens and 4X optical zoom using hybrid (HYD) detectors. The acquired images were analyzed using Fiji software and the quantitation was done manually by counting the γH2Ax puncta in each cell.

### Small cell number input Western Blot

Briefly, 5000 mouse LT-HSCs were isolated from middle-aged SCD and non-SCD mice as described above while sorting directly into 20µL of PBS with 2% FBS containing protease and phosphatase inhibitors (Invitrogen, Carlsbad California). Next, an appropriate volume of 4X LDS sample buffer containing reducing agent was added to each sample such that the final concentration was 1X, and cell lysis and protein linearization were achieved by heating samples to 95°C for 5 minutes. Polyacrylamide gradient gel electrophoresis, immunoblotting, and analysis were performed as previously described(*81, 82*) using primary antibodies against RPL21 (A305-031A, Bethyl Laboratories, Montgomery, TX), H2A (D6O3A, Cell Signaling Technologies, Danvers, MA), and β-actin (13E5, Cell Signaling Technologies, Danvers, MA) at a 1:1000 dilution and anti-rabbit HRP conjugated secondary antibody (#7074, Cell Signaling Technologies, Danvers, MA) at 1:10,000 dilution.

### Extreme Limiting Dilution Analysis

Extreme limiting dilution analysis and estimation of long-term blood repopulating cells was performed using web based software as previously described(*83*), and this webtool also subjects the resultant data to tests for goodness of fit and heterogeneity. The likelihood ratio test of single model hit on our data from young SCD and non-SCD WBM resulted in χ^2^=0.0677, P=0.795, df=1 and χ^2^=2.35, P= 0.125, df=1 for the data from aged SCD and non-SCD WBM, indicating that we cannot reject the single-hit model.

### SA-β-Galactosidase activity by staining of mouse LT-HSC cytospin preparations

Mouse LT-HSC SA-β-Gal activity was also assayed and confirmed by using FACS to isolate LT-HSCs from middle-aged SCD and non-SCD mice which were then mounted on microscopy slides by cytospin at 500 g for five minutes at room temperature. LT-HSCs were then stained for SA-β-Gal activity using the Senescence Detection Kit (Millipore Sigma, Burlington, MA) per manufacturer’s instructions. SA-β-Gal^+^ cells were then assessed visually by bright field microscopy of 10 random fields of view per biological replicate

### Bulk RNA-sequencing of mouse LT-HSCs and human HSPCs

Mouse bone marrow LT-HSCs or human bone marrow HSPCs (CD45^+^Lineage^-^CD34^+^CD38^-^) were collected by FACS directly into the provided RNA lysis buffer for isolation of total RNA (RNeasy Micro kit; QIAGEN, Germantown, MD). Cells were collected from either SCD or non-SCD littermates (*i.e.* Townes mice) or cryopreserved SCD and non-SCD patient bone marrow samples. The Ovation RNA Seq System V2 kit (Tecan, Männedorf, Switzerland) was used for library preparation for mouse cells, while the SMART-Seq v4 Ultra Low Input RNA Kit (Takara Bio, Kusatsu, Japan) was used for human cells. 150 bp paired-end sequencing was performed on the Illumina NovaSeq 6000, targeting an average of 50 million reads/sample by the Vanderbilt Technologies for Advanced Genomics Genomics core laboratory (Vanderbilt University Medical Center, Nashville, TN).

### Gene expression data analysis for bulk RNA-Sequencing

Human and mouse datasets were analyzed separately via similar strategies. For both mouse and human bulk RNA-Seq datasets, technical quality of reads was checked with FastQC (version 0.12.0)(*84*) before and after read trimming. Reads were trimmed and filtered with fastp (version 0.23.4)(*85*) to remove adapter sequence, low-quality bases near the ends of reads, and to remove reads with fewer than 40 high-quality bases. Library quality was assessed with BBMap (version 38.86)(*86*) and RSeQC (version 3.0.1)(*87*). Filtered reads were mapped to their respective genome with STAR (version 2.7.11)(*88*). For human, the T2T-CHM13v2.0 genome assembly with annotation from Ensembl (release 110.1)(*89*) was used, and for mouse the GRCm39 genome assembly with annotation from Gencode (version M35)(*90*) was used. Gene-level counts were obtained with featureCounts from Subread (version 2.0.5)(*91*) and transcript-level expression was quantified with RSEM (version 1.3.3)(*92*). Up to 20 alignments for a given read were output from STAR, but multi-mapping reads were not counted by featureCounts.

Downstream analysis was performed primarily in R (version 4.3.2)(*93*) with packages from the Bioconductor repository (version 3.18)(*94*). For both mouse and human datasets, features with 10 or more raw counts in any sample within the respective dataset were retained; these were further filtered to exclude gene types which have not been reported to exhibit polyadenylation, leaving 14,012 coding genes for mouse (18,245 total expressed features), and 15,699 coding genes for human (19,320 total expressed features). For human data, due to considerable sequence identity resulting in ambiguous mapping with short-read sequencing, read counts for the paralogous pairs of gamma globin (HBG1, HBG2) and alpha globin (HBA1, HBA2) genes were summed and represented by meta-features, HBG1_HBG2 and HBA1_HBA2, respectively. Exploratory analysis and visualization (PCA, clustering, sample-sample correlation, expression heatmaps) was performed using normalized counts after variance-stabilizing transformation (VST) as implemented in DESeq2(*95*), or in transcript per million (TPM) estimates from RSEM. The factoextra R package(*96*) was used for PCA plots with confidence ellipses, with the ellipses representing the 95% confidence interval for the indicated PCs and sample groups based on a normal distribution. Differential gene expression analysis was facilitated by DESeq2, using raw counts as input. The resulting log_2_ fold-change estimates were moderated to reduce the apparent effect size of genes with low or highly variable expression with the apeglm fold-change shrinkage method(*97*).

For the mouse dataset, one sample with low library complexity was identified as an outlier from PCA and removed, leaving n=4 SCD and n=5 non-SCD samples for differential expression. For the human dataset, biological sex was included as a blocking factor in the design formula for differential expression (design ∼ sex + Condition) and DE analysis was run first with all samples (n=7 SCD, n = 4 non-SCD), and then on a subset of samples (n=3 SCD and n=3 non-SCD) found to separate by condition on PC3 in the PCA with all 11 samples. Genes with Benjamini-Hochberg FDR-adjusted(*98*) p-values <0.05 and |log_2_ fold-change| ≥0.5 were considered as DEGs except for the human analysis with all 11 samples, where only the FDR p-value threshold was applied. Functional analysis for DEGs was performed using GOSeq(*99*) to test for over-representation of pathways and gene sets assembled from KEGG(*100*), Gene Ontology(*101*), MSigDB(*102*), CellAge(*103*), SenNet(*104*), and SenMayo(*105*). Functional terms and pathways with FDR-adjusted enrichment p-values < 0.05 were considered significant. Gene Ontology term network representations were created in Cytoscape (version 3.10.2)(*106*) with enrichment results from GOSeq. Where necessary, gene symbols were mapped between mouse and human using homology information from HGNC and MGI(*107, 108*).

### Sample preparation for mouse CITE-Seq

SCD and non-SCD mouse bone marrow samples from two and six-month old animals (n =4/group) were enriched for c-Kit^+^ cells (*i.e.* total HSPCs) via magnetic enrichment using anti-CD117 microbeads and an autoMACs magnetic cell separator (Miltenyi Biotec, Carlsbad, CA) per manufacturer’s instructions. c-Kit-enriched samples were prepared for CITE-seq by staining with a panel of Biolegend TotalSeq-A antibodies (Ly-6A, CD150, CD48, CD16/CD32, CD105, CD41, and CD71). Towards this, samples were incubated at 4°C for 30 minutes in 50 µL of Cell Staining buffer (Biolegend, Catalog# 42020) containing TotalSeq-A antibodies (0.02 µg/mL per antibody), and then washed and resuspended in 2%FBS (1.5×10^6^/mL) for downstream single-cell RNA sequencing. Labeled cells were processed for single cell RNA sequencing using the Chromium Single Cell 3’ Reagent Kit v3 with a Chromium Controller (10x Genomics) and libraries were prepared for sequencing according to the manufacturer’s instructions, yielding libraries for both gene expression and antibody-derived tags (ADTs). Two-month old animals were processed for CITE-Seq and 10x cell capture in one batch, while six-month animals were processed in another, due to limited lanes on the 10x microfluidic chip. Sequencing for each library type was performed with all samples together on an Illumina NovaSeq 6000.

### Data processing and analysis for single cell sequencing

Single cell RNA-Sequencing FASTQ files were aligned to the mouse genome (GRC m39 with GENCODE annotation M33) and quantified with the STAR aligner’s (v2.7.11) STARsolo workflow(*109*) (parameters for read trimming and filtering and cell filtering: “--clipAdapterType CellRanger4 --outFilterScoreMin 30 --soloCellFilter EmptyDrops_CR 3000 0.99 10 45000 90000 500 0.01 20000 0.01”). Cell surface protein tag data was counted with CITE-seq-Count (v.1.4.4, https://github.com/Hoohm/CITE-seq-Count) using the tag sequences provided by Biolegend. Read counts for both assays were first read into R and assembled into one Seurat object per sample (Seurat v5)(*110*). Gene expression was normalized with SCTransform v2(*111*), with each sample processed separately and merged post-normalization. Antibody-derived tag data was normalized with ADTnorm(*112*). Cells found in both gene and cell surface protein assays, having fewer than 10% mitochondria-derived reads, and with expression of at least 500 genes were taken forward. Putative doublets/multiplets (more than one cell captured in a single droplet) were removed through an ensemble approach based on gene expression including: (1) cells that were outliers for total read counts or total expressed genes (> 3 MAD for either), (2) cells that were outliers for total read counts or total expressed genes and having a score from scDblFinder(*113*) > 0.9, and (3) cells having both scDblFinder score > 0.9 and called as a doublet by DoubletFinder(*114*). Genes expressed in 50 or fewer cells across the filtered dataset were removed. An ensemble approach was also implemented to filter cells out extreme outliers based on ADT expression level, which could represent empty droplets (much lower than most values) or multiplets (multiple times higher than most values). Briefly, cells were removed if they met any three of four criteria: (1) > 10 MAD on the right tail for any antibody, (2) ≥ limit determined by histogram-based outlier detection in the R package “bigutilsr”(*115*) with a maximum outlier detection fraction of 0.01 and 100 bootstrap iterations, (3) called as an outlier by the “adjusted outlyingness” method as implemented in the R package “univOutl”(*116*), and (4) called as an outlier by the Robust Kernel-based Outlier Factor with bootstrap cutoff determination as implemented in the R package “OutlierDetection”(*117*). An average of approximately 4858 cells per sample remained in the final dataset after cell filtering.

Unsupervised cell clustering was performed on SCT-normalized gene expression using the Louvain algorithm as implemented in Seurat. Clustering parameters were swept over ranges to identify sub-ranges leading to stable cluster membership and cluster count, from which the final parameter values were chosen. Harmony(*118*) was applied to reduce the influence of preparation batch on dimension reduction and projection into UMAP-space. Cell cycle inference was performed with Seurat’s cell cycle scoring function after converting gene symbols for the provided human cell cycle markers to mouse homologs.

Cell type annotation was performed at a single-cell-level, as summarized in Extended Data Fig. 6a. First, well-characterized reference single-cell datasets were obtained pre-processed from data repositories, checked for quality, and normalized with SCTranform v2. The following reference datasets were used: mouse HSPC profiles from Nestorowa *et al.* (obtained from the R package scRNAseq), bone marrow and spleen profiles assayed with 10x scRNA-Seq and SMART-Seq from Tabula Muris (obtained from ExperimentHub), and single-cell data from the Immunological Genome Project (obtained from CellDex)(*119–123*). Scores for cell types were assigned to each cell with SingleR for each reference dataset and summarized by finding the highest-scoring type for a given cell within each reference. Next, in an orthogonal approach, sets of cell type marker genes for normal blood and bone marrow curated from literature and included with the R package *scCATCH* were used as input gene sets for single-cell gene set activity scoring with AUCell(*124, 125*). The maximum scoring set was identified for each cell, with the few ties broken manually by checking expression of other common secondary markers for the cells in question. If the maximum AUCell score was < 0.6, cell type was set to “undetermined” to indicate a lack of high-confidence matches among the queried signatures from scCATCH. Finally, scores from across the two approaches and reference datasets were integrated to assign a cell type in a tiered approach: (1) the score differences (Δ_score_) between the best-scoring SingleR cell type from Nestorowa *et al.* and the best-scoring types from the other SingleR reference sets and AUCell were found for each cell, facilitated by the SingleR and AUCell scores sharing a 0-1 scale. (2) The means of Δ_score_ for each cell were calculated. (3) For a given cell, if mean[Δ_score_] ≥ −0.25, the best scoring type from Nestorowa *et al.* was assigned. [Δ_score_] < −0.25 indicated that other reference data sources scored substantially higher and thus the cell type corresponding to the highest alternative score from the other results was assigned. This threshold was selected after considering the distribution of mean[Δ_score_]. Cell types with very low representation in the final annotation set (< 200 cells) were set to type “other”, which represented 0.55% of the cells in the dataset. See Extended Data Fig. 6c for the number of cells of each type that were annotated based on each reference dataset.

Differential expression analysis was performed using a pseudobulk strategy with DESeq2 for comparisons of SCD versus non-SCD within cell types when not considering finer cell clustering structure. To identify DEGs within and between distinct clusters of a given cell type, particularly LT-HSCs for which not all biological replicate samples were represented in every cluster, MAST(*126*) was utilized for testing. For both strategies, significant DEGs were called based on FDR-adjusted p-value < 0.05 and |fold-change| ≥ 1.5. Functional enrichment was performed for DEG results using gene sets from the mouse version of MSigDB and GOSeq, as otherwise described under “*Gene expression data analysis for bulk RNA-Seq”*.

### Time-lapse live cell imaging

Live-cell imaging experiments were done in IMDM supplemented with 20% BIT (Stem Cell Technologies), 1% P/S, 2mM L-Glutamine, 50uM 2-mercaptoethanol, 100ng/mL mouse SCF and 100ng/mL TPO. All images were acquired using a Nikon Ti-Eclipse 2 microscope with a motorized stage, a 10.2 Megapixel sCMOS camera (Kinetix, Photometrics) with a 6.5 µm x 6.5 µm pixel area, a Spectra III fluorescent light source (Lumencor) and a transmitted light LED from Nikon. Experiments were done at 37 °C, 5% O2, and 5% CO2 using a manual three-gas mixer (Okolab) and custom-made microscope enclosures with a heating unit controlled by Oko-touch (Okolab). Custom-made 3D-printed stagetop incubators and humidifiers were used to prevent evaporation. Fluorescent images were acquired using custom-made optimized filters from Chroma or Semrock: BV480 (436/20; 455LP; 470/24), a488 (488/10; 495LP; 532/18), PE (555/28; 555LP; 575/25), APC (620/30; 645LP; 680/13) to detect anti-CD16/32-BV480, anti-Sca1-a488, anti-CD41-PE, anti-CD71-APC, respectively. Bright-field and fluorescent images were acquired every 30 minutes and light intensities and exposure were selected to minimize phototoxicity. Images were acquired with a 10x CFI Plan Apochromat λ objective (NA 0.45). Single-cell tracking and image quantification were done with self-written software as described(*127, 128*). Images with 3200×3200 pixel resolution were acquired as 16-bit .tiff and linearly transformed to 8-bit .png files using optimized white points. Shading of fluorescence images was corrected using BaSiC(*129*). Image quantification was done using fastER(*130*) using brightfield channel segmentation and dilation of labeling masks (Settings: dilation 8) to ensure reliable quantification of the entire cell. Tracking and quantification of fluorescence channels were done as described(*127, 128*) and analyzed using Matlab 2023a (Mathworks).

Criteria for excluding individual data points were determined in advance. For Time-Lapse Imaging Experiments: Time points of single-cell time series were curated and cross-validated by another scientist. Segmentation masks resulting from over- and under-segmentation were excluded. Also, cells touching the image border, cell clumps, cells touching fluorescence debris, and loss of single-cell identities during tracking were excluded.

Time-lapse images were acquired using commercially available software (NIS-Elements 5.3.2). Image segmentation, single-cell tracking and fluorescence quantification analysis tools used in this study are published and open-sourced (see https://doi.org/10.1038/nbt.3626) and (https://academic.oup.com/bioinformatics/article/33/13/2020/3045025). The code for dimensionality reduction using Uniform Manifold Approximation and Projection (UMAP) is published and open-sourced(*131*).

## Supplementary Figures

**Supplementary Data Figure 1.**
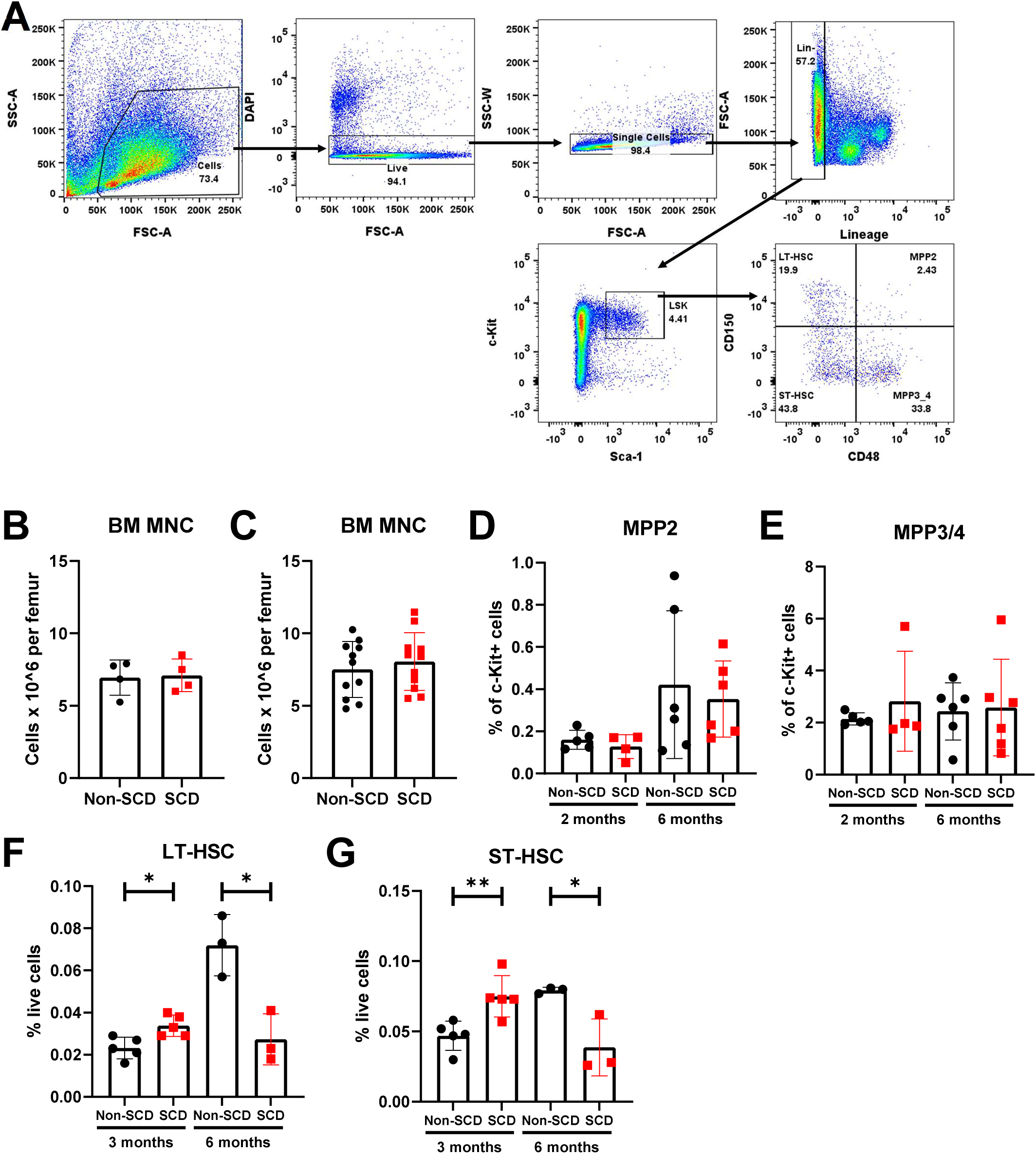
Mouse HSPC gating strategy, total bone marrow cellularity, MPP frequencies and HSCs in Berkeley mouse BM. **A**, Representative flow cytometry gating strategy for phenotypic mouse LT-HSCs, ST-HSCs, MPP2s, and MPP3/MPP4s. **B**, Quantification of bone marrow cellularity per leg in young SCD and non-SCD mice (n=4 each). **C**, Quantification of bone marrow cellularity per femur in middle aged SCD and non-SCD mice (n=11 each). **D**, Quantification of MPP2 frequencies in young (left) and middle-aged (right) SCD and non-SCD mice (n=4-6 each). **E**, Quantification of MPP3/MPP4 frequencies in young (left) and middle-aged (right) SCD and non-SCD mice (n=4-5 each). **F**, Quantification of LT-HSC frequencies in young (3 months old, left) and middle-aged (right) Berkley SCD and non-SCD mice (n=3-5 each). **G**, Quantification of ST-HSC frequencies in young (3 months old, left) and middle-aged (right) Berkley SCD and non-SCD mice (n=3-5 each).

**Supplementary Data Figure 2.**
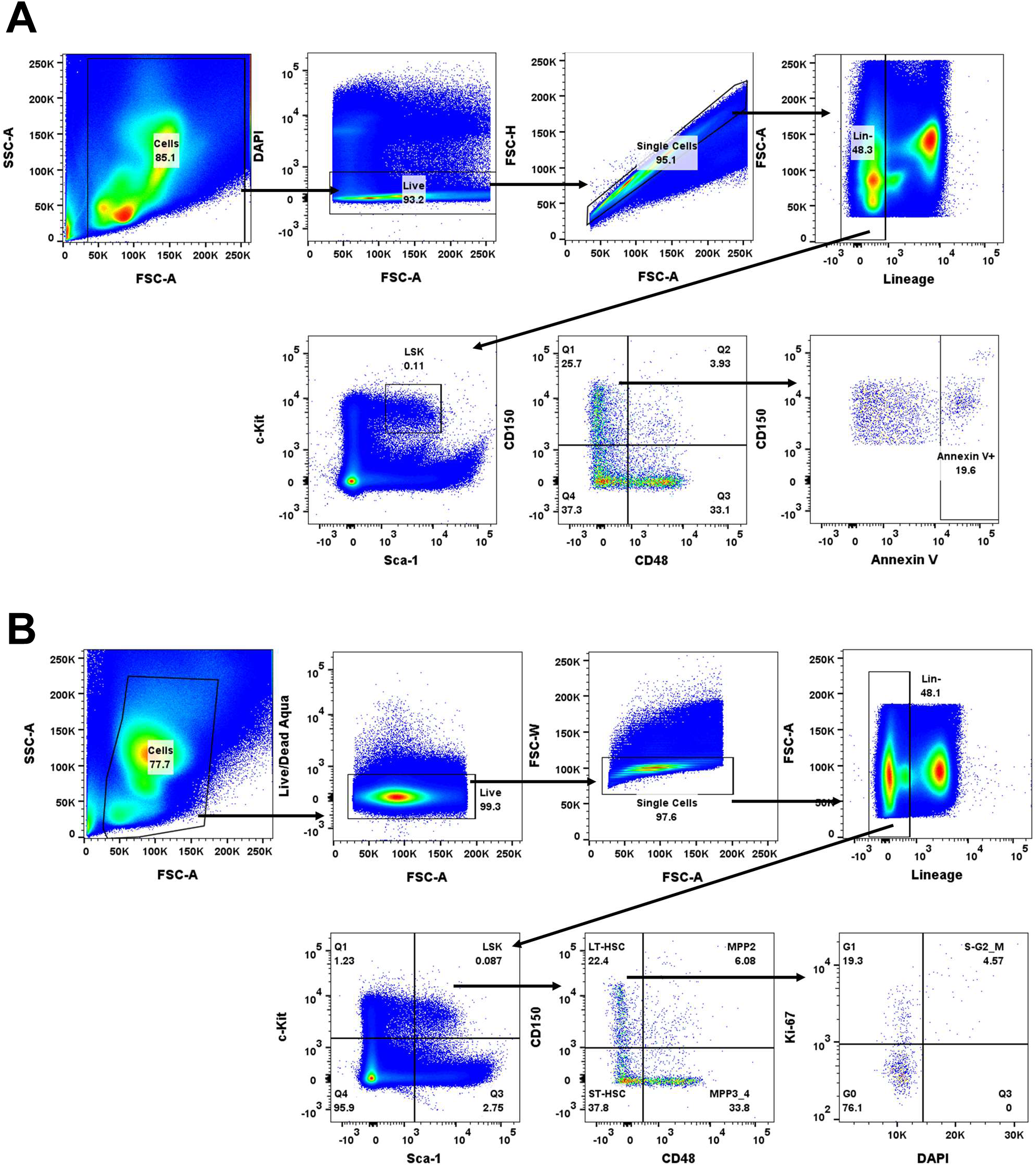
Mouse LT-HSC gating strategy for cell cycle and apoptosis **A,** Representative flow cytometry gating strategy for identification of apoptotic (Annexin V^+^) LT-HSCs in SCD and non-SCD mice. **B,** Representative flow cytometry gating strategy for identification of LT-HSCs in each phase of cell cycle (G0, G1, or S/G2-M) by intracellular staining for the proliferation protein, Ki-67, and cellular DNA content by DAPI.

**Supplementary Data Figure 3.**
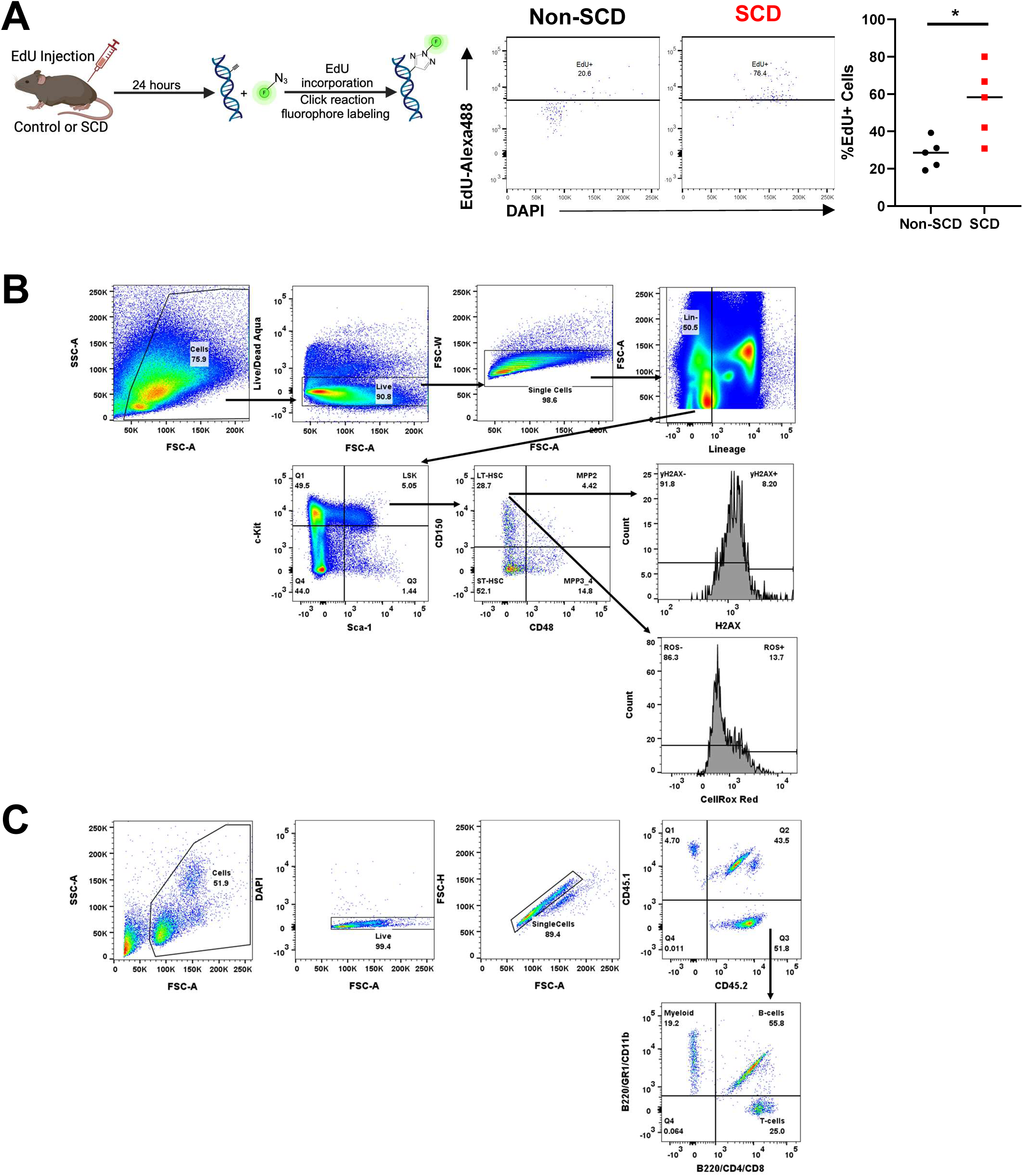
Increased in vivo EdU uptake of LT-HSCs in SCD mice and representative gating strategies for DNA damage, ROS, and peripheral blood repopulation post-transplant. **A**, Experimental schematic, representative flow cytometry plots (left), and quantification (right) of %EdU^+^ bone marrow LT-HSCs in middle-aged SCD and non-SCD mice (n=5 each). **B,** Representative flow cytometry gating for γH2AX^+^ (pH2AX) LT-HSCs and LT-HSCs with an increased amount of total ROS, as reflected by CellRox Red staining. **c,** Representative flow cytometry gating for CD45.2^+^ PB and PB lineages.

**Supplementary Data Figure 4.**
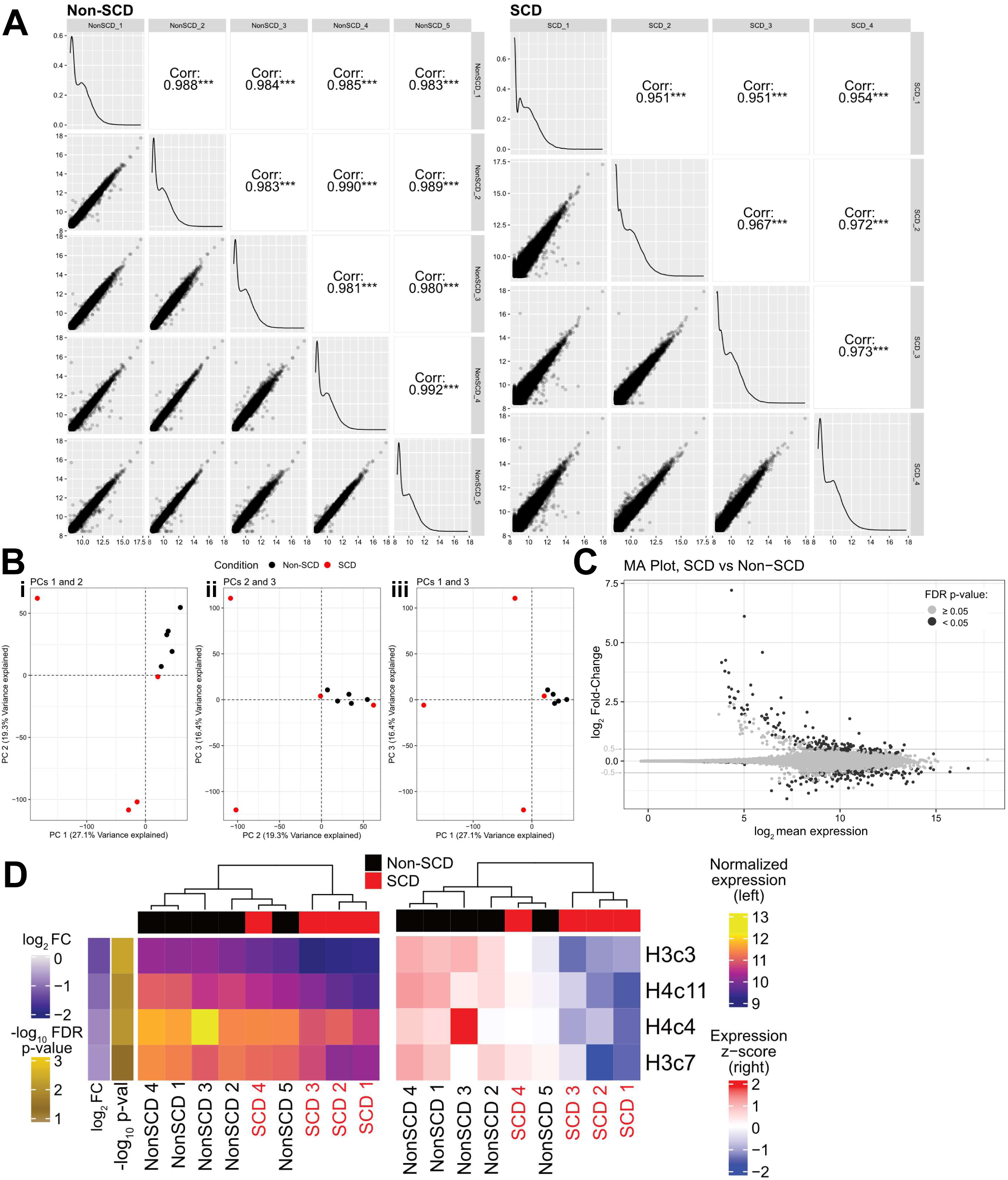
Additional analyses of bulk RNA-Sequencing of LT-HSCs from SCD and non-SCD mice. **A**, Pairwise Pearson correlation between biological replicate samples across all 18,254 features expressed in dataset shows high overall correspondence between replicates. Intra-group correlations are higher for non-SCD than SCD mice, corroborating higher variability across SCD samples also seen via PCA (Fig. 3). **B**, A subset of SCD samples exhibits substantially greater variability than non-SCD in each of the first three principal components. (**i**) PC1 versus PC2, (**ii**) PC2 versus PC3, and (**iii**) PC1 versus PC3. SCD (n=4) and non-SCD (n=5) mice. **C**, MA plot illustrating the relationship between average expression (log_2_ scale) and fold-change between SCD and non-SCD mice-derived LT-HSCs (log_2_ scale) for all expressed features after fold-change shrinkage with apeGLM through DESeq2 to moderate fold-changes for genes with very low or highly variable expression. Gray horizontal bars shows the fold-change threshold which was used as part of the DE significance criteria and the color of the points indicates if the FDR-adjusted p-value was <0.05 (black) or not significant (gray). **D**, Heatmaps of histone subunit genes with significantly altered expression in SCD mouse LT-HSCs. Left: VST-normalized expression, right: z-scores across rows to highlight the trends in expression for each gene across samples. Samples are ordered by hierarchical clustering and rows are ordered by fold-change. Adjusted p-values are also indicated in the left side bar, with brighter gold indicating lower p-values; all the genes shown had FDR p-value <0.05.

**Supplementary Data Figure 5.**
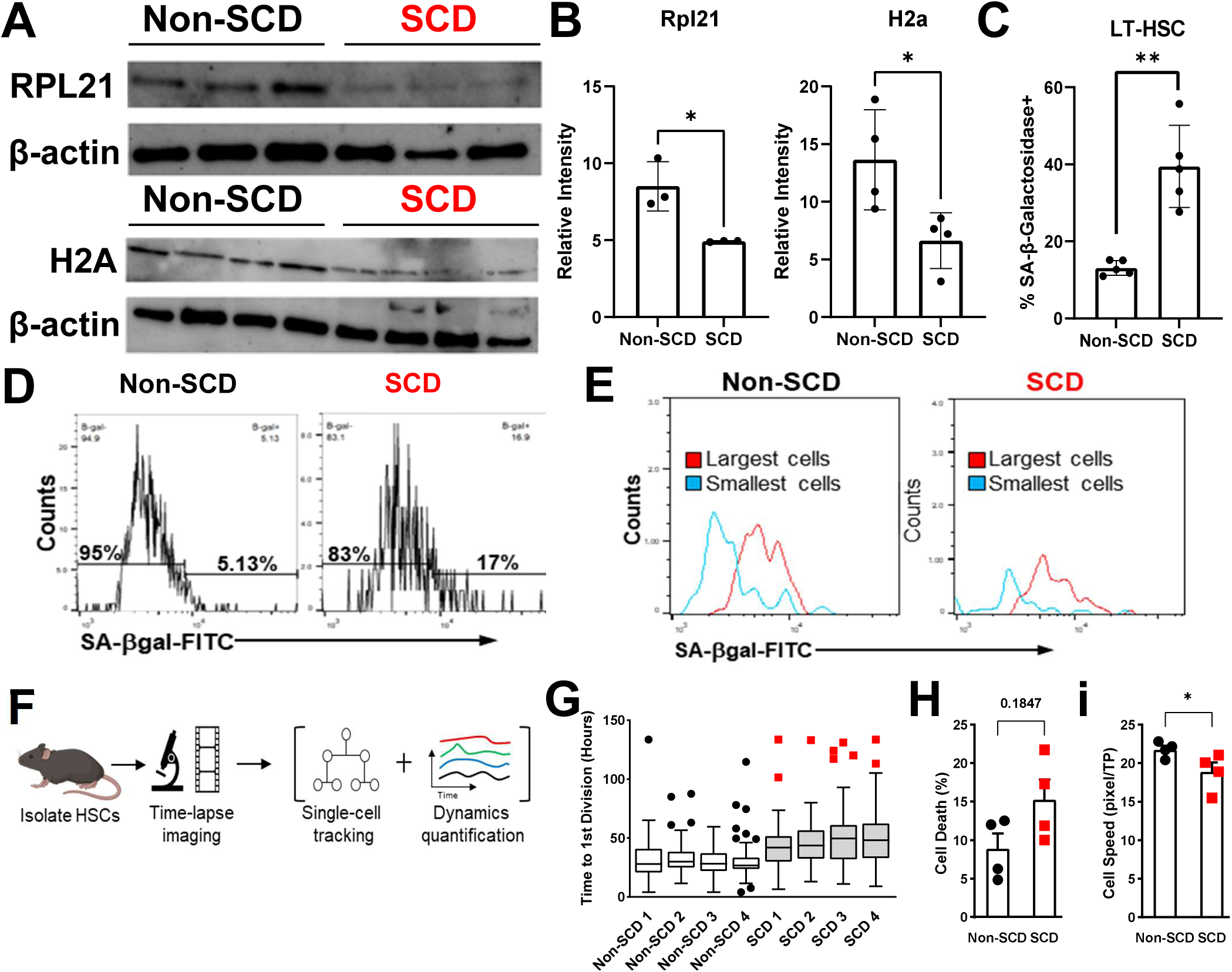
LT-HSCs from middle-aged SCD mice display features of senescence. **A**, Small cell input western blot for RPL21 (top, n=3 for non-SCD; n=3 for SCD) and H2A (bottom, n=4 for non-SCD; n=4 for SCD) in LT-HSCs isolated from middle-aged SCD and non-SCD mice. Each lane contains protein extracted from 5000 LT-HSCs. **B**, Densitometric quantification of RPL21 (left) or H2A (right) band signal intensity from (a) normalized to corresponding respective β-actin band signal intensity. **C**, %SA-β-Gal+ LT-HSCs in middle-aged SCD and non-SCD mice as determined by cytospin and SA-β-Gal staining on microscopy slides (n=5 each). **D**, Representative flow cytometry histograms showing SA-β-Gal staining in LT-HSC. **E**, Representative flow cytometry histograms demonstrating that LT-HSCs tend to be larger in middle-aged SCD mice, relative to non-SCD mice, and that SA-β-Gal+ LT-HSCs are larger than SA-β-Gal-LT-HSCs. **F**, Schematic of live cell imaging experiments. Quantification of LT-HSC time to first division (**G**), cell death (**H**), and cellular motility (**I**) at the end of the duration of live cell imaging of individual LT-HSCs isolated from middle-aged SCD and non-SCD (n=4 each).

**Supplementary Data Figure 6.**
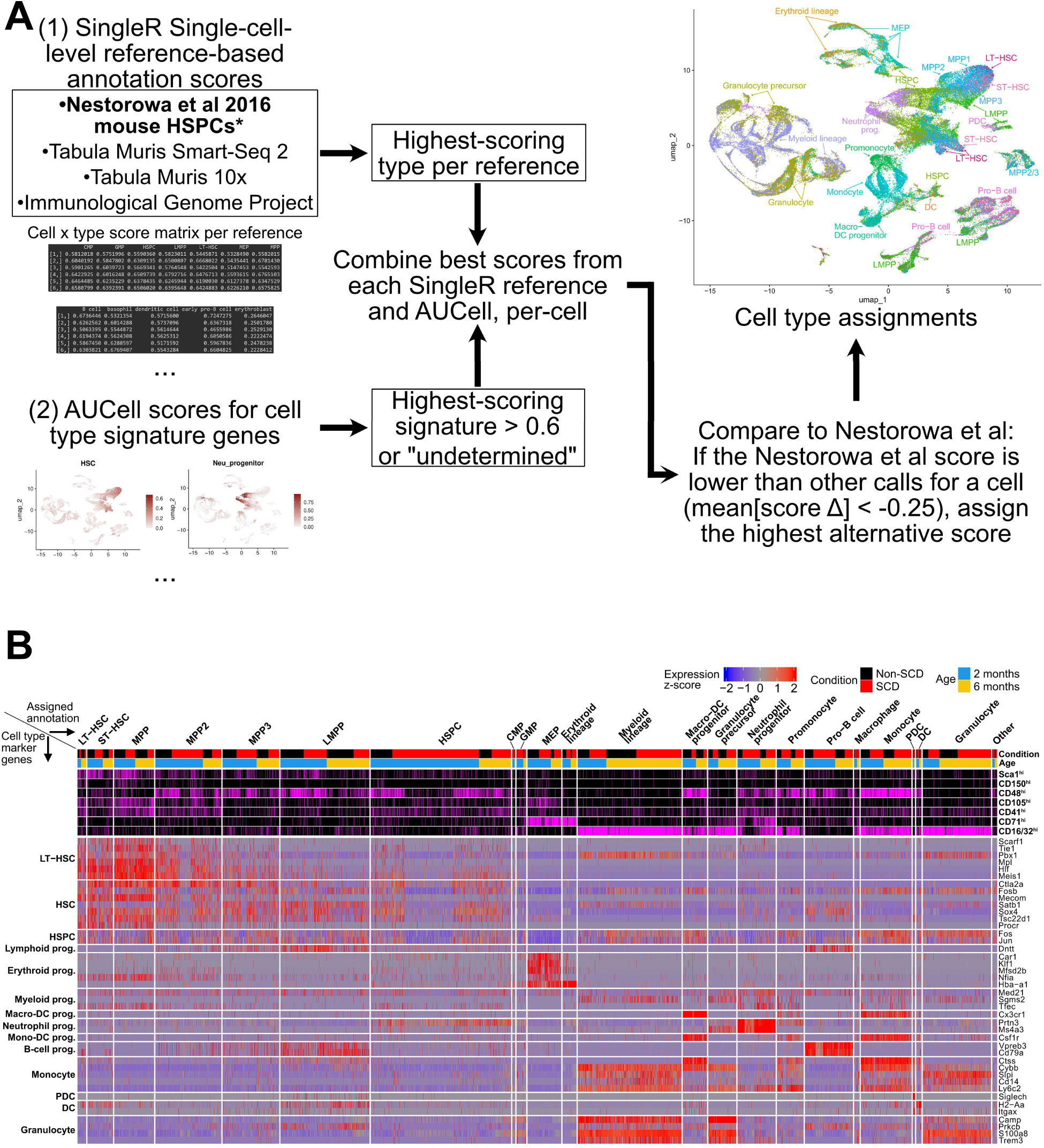
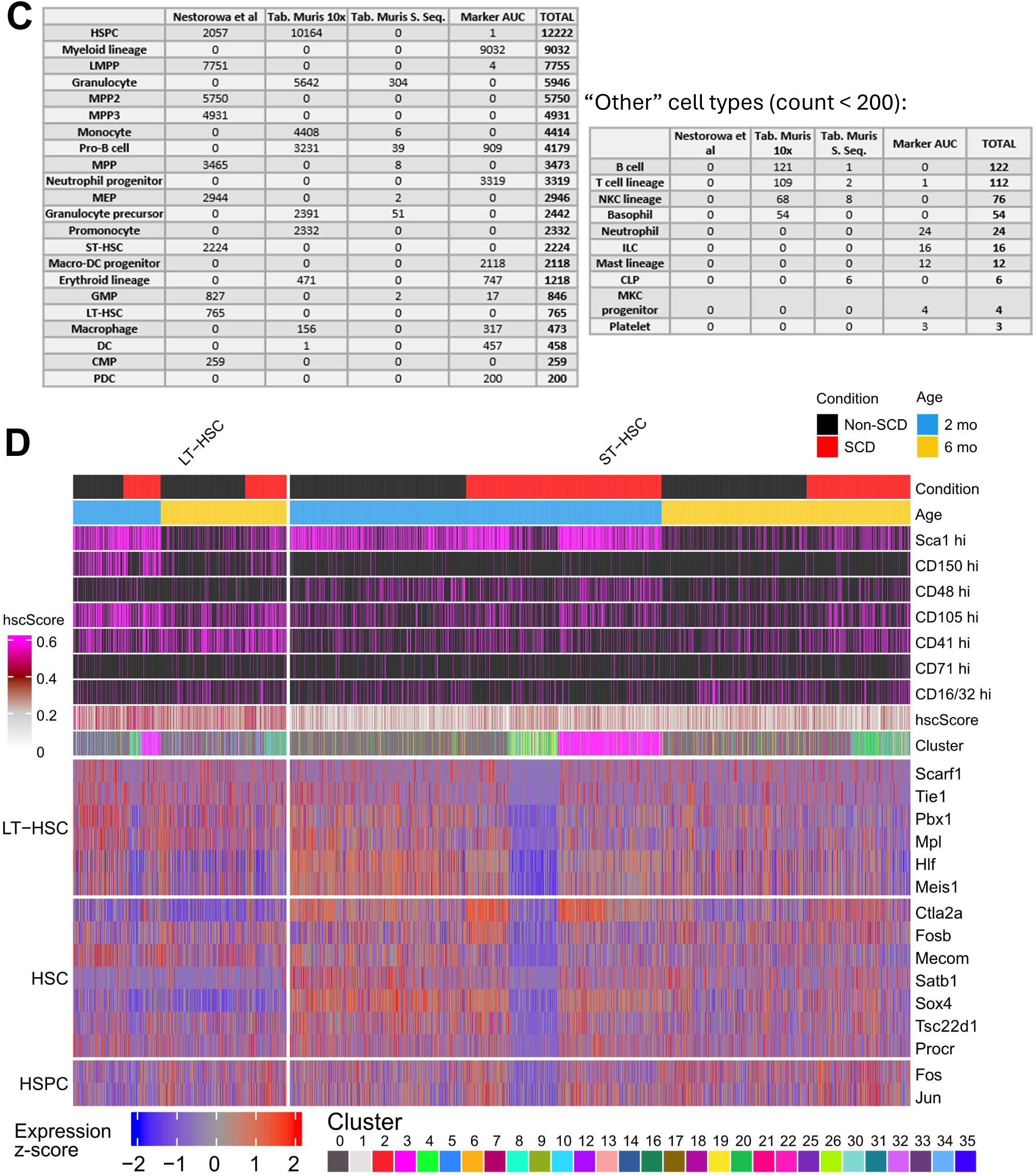
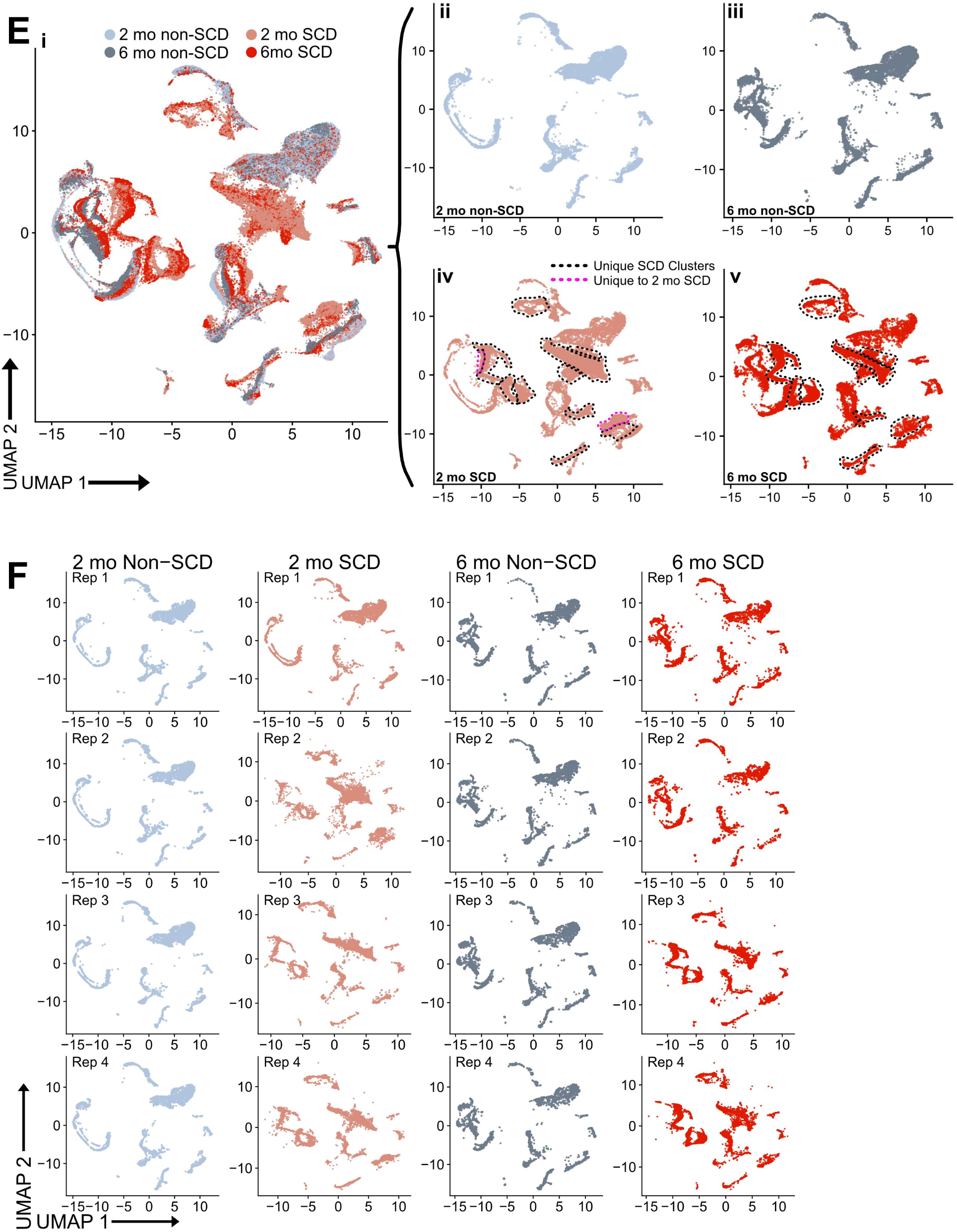
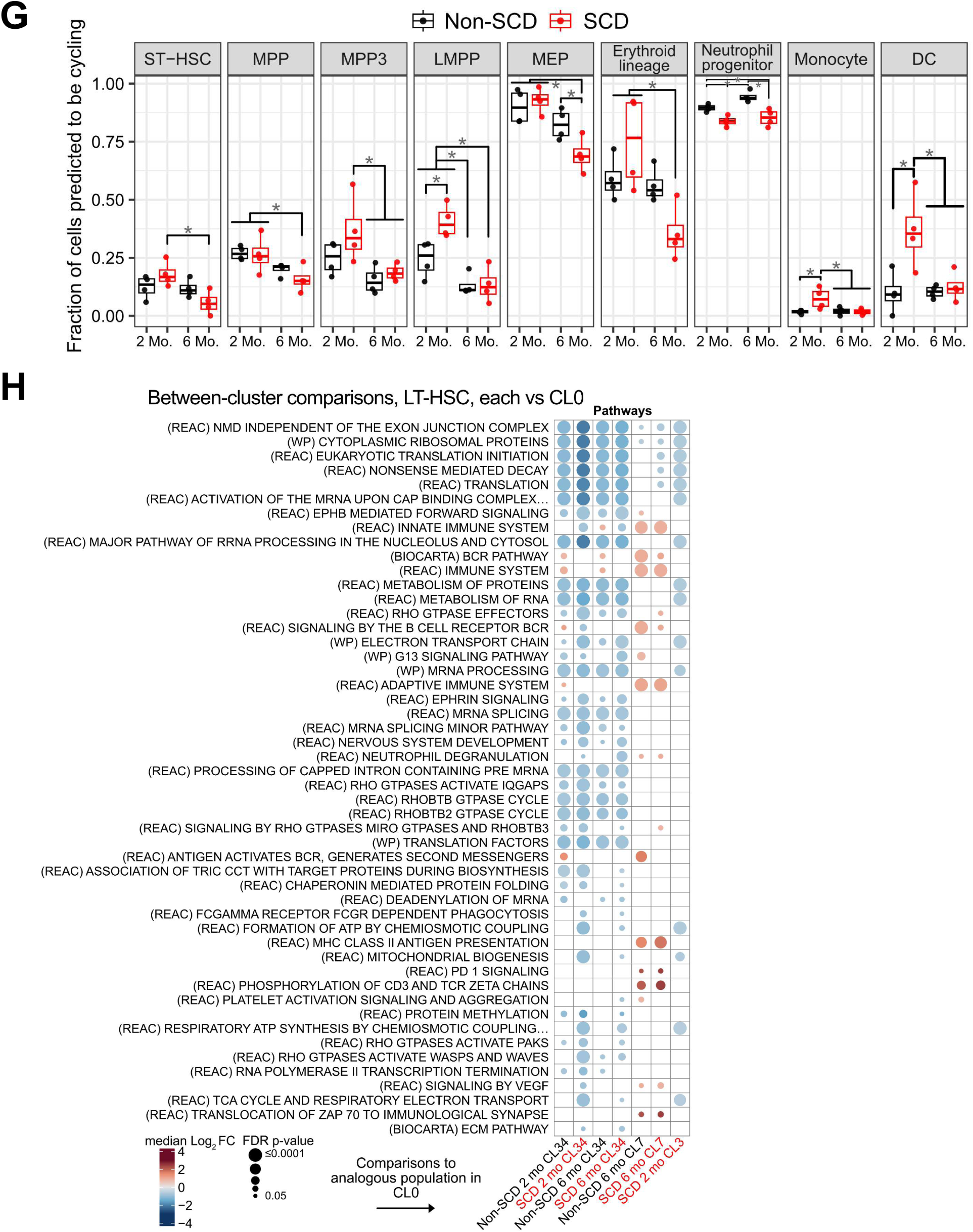
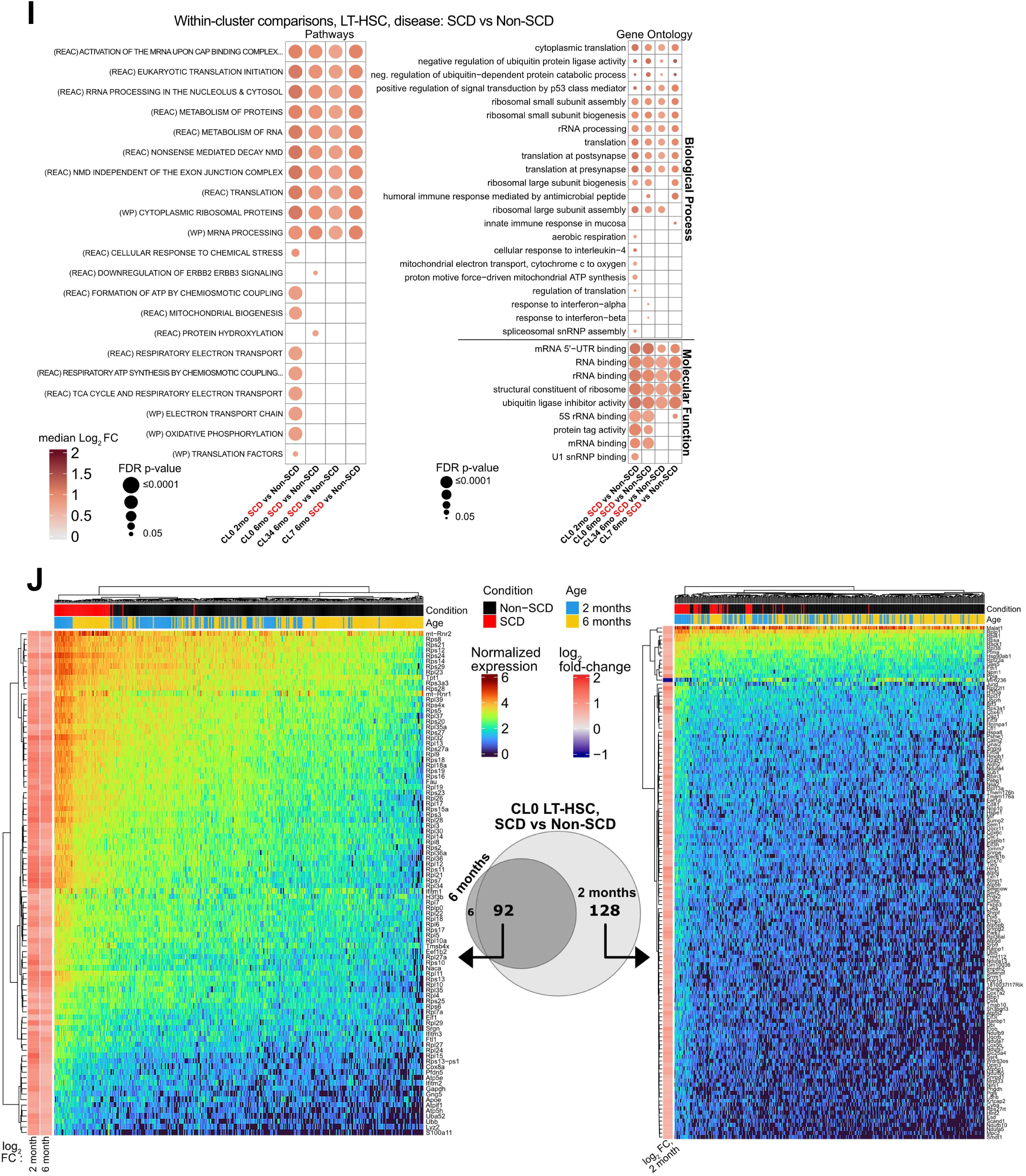
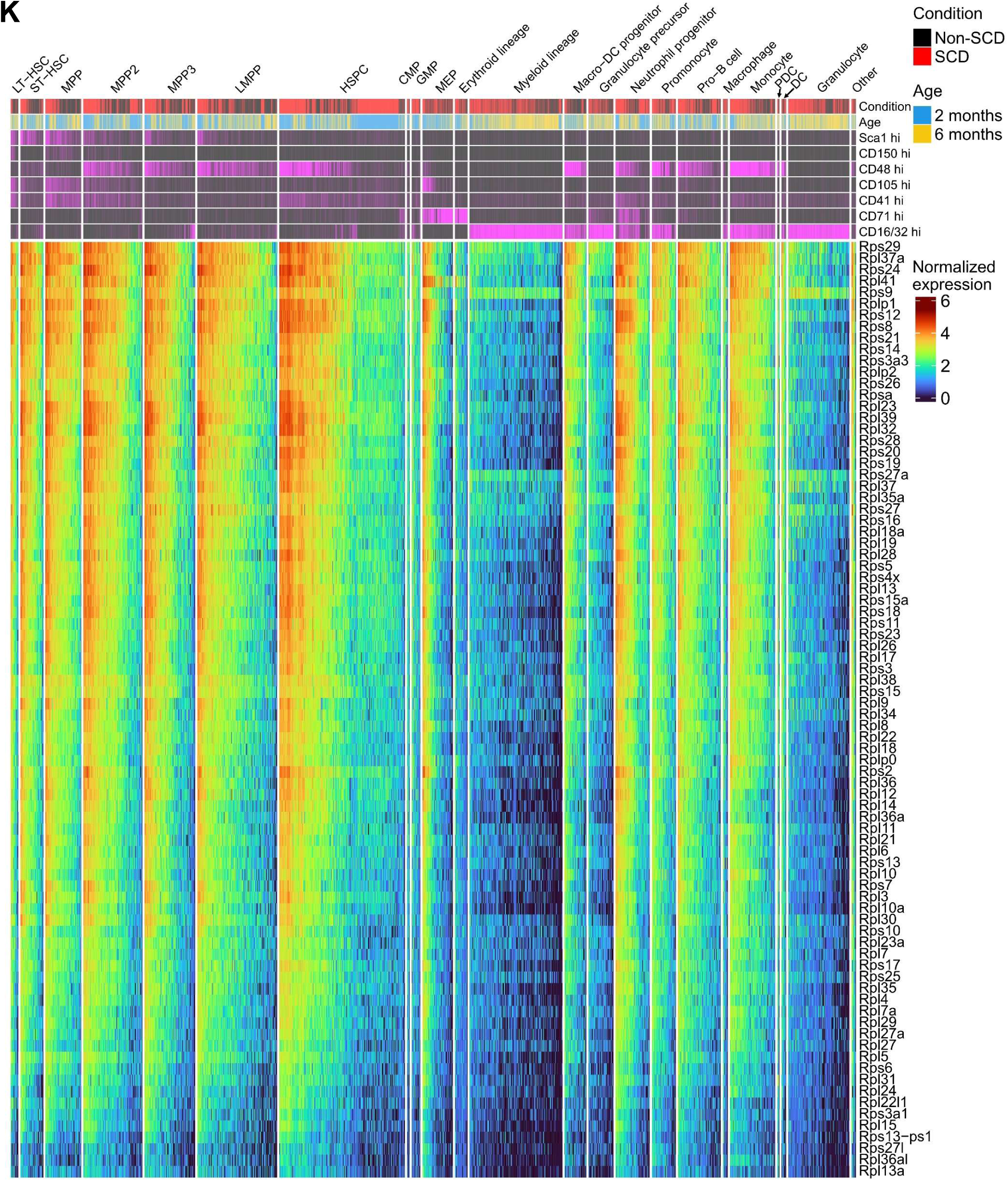
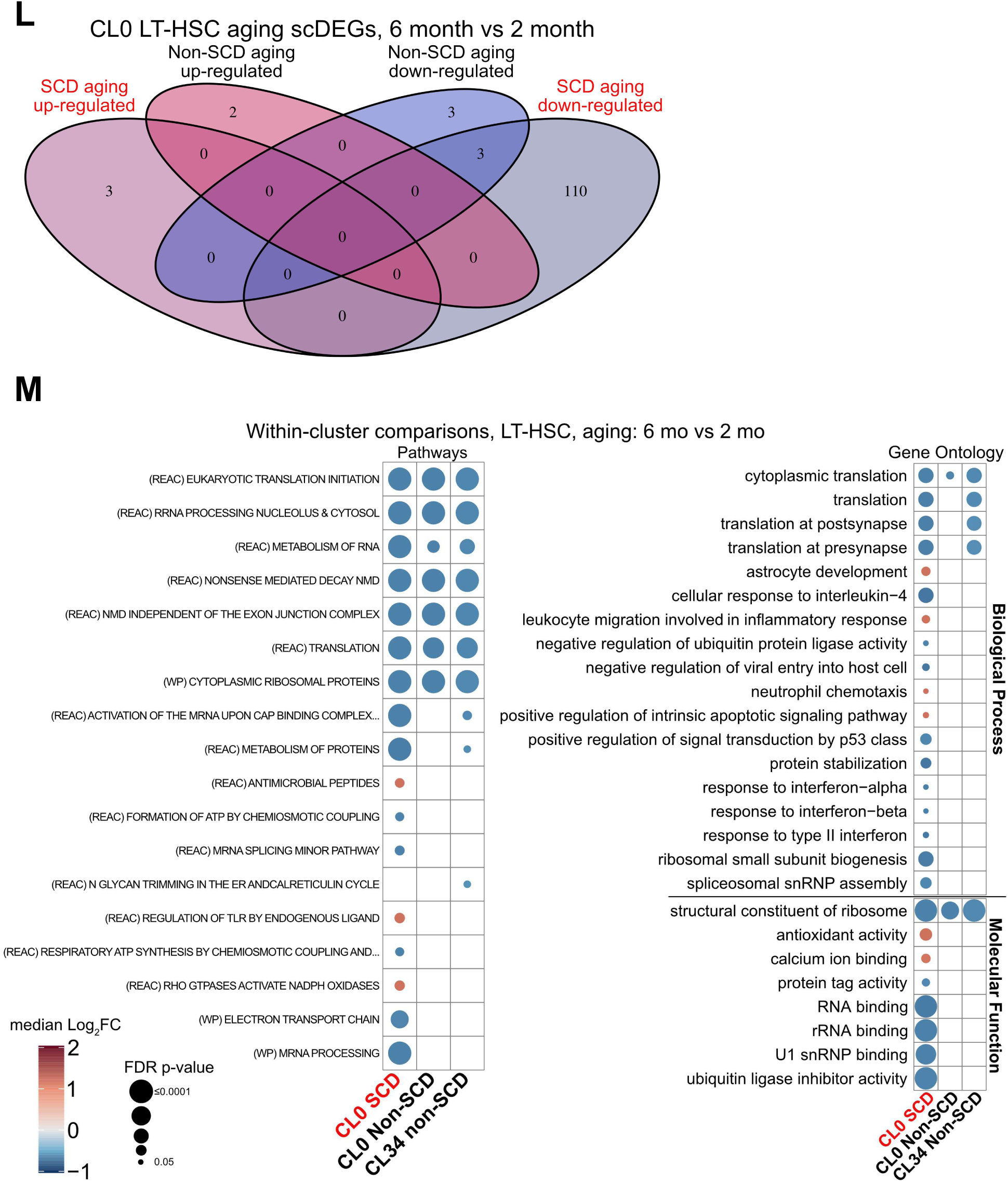
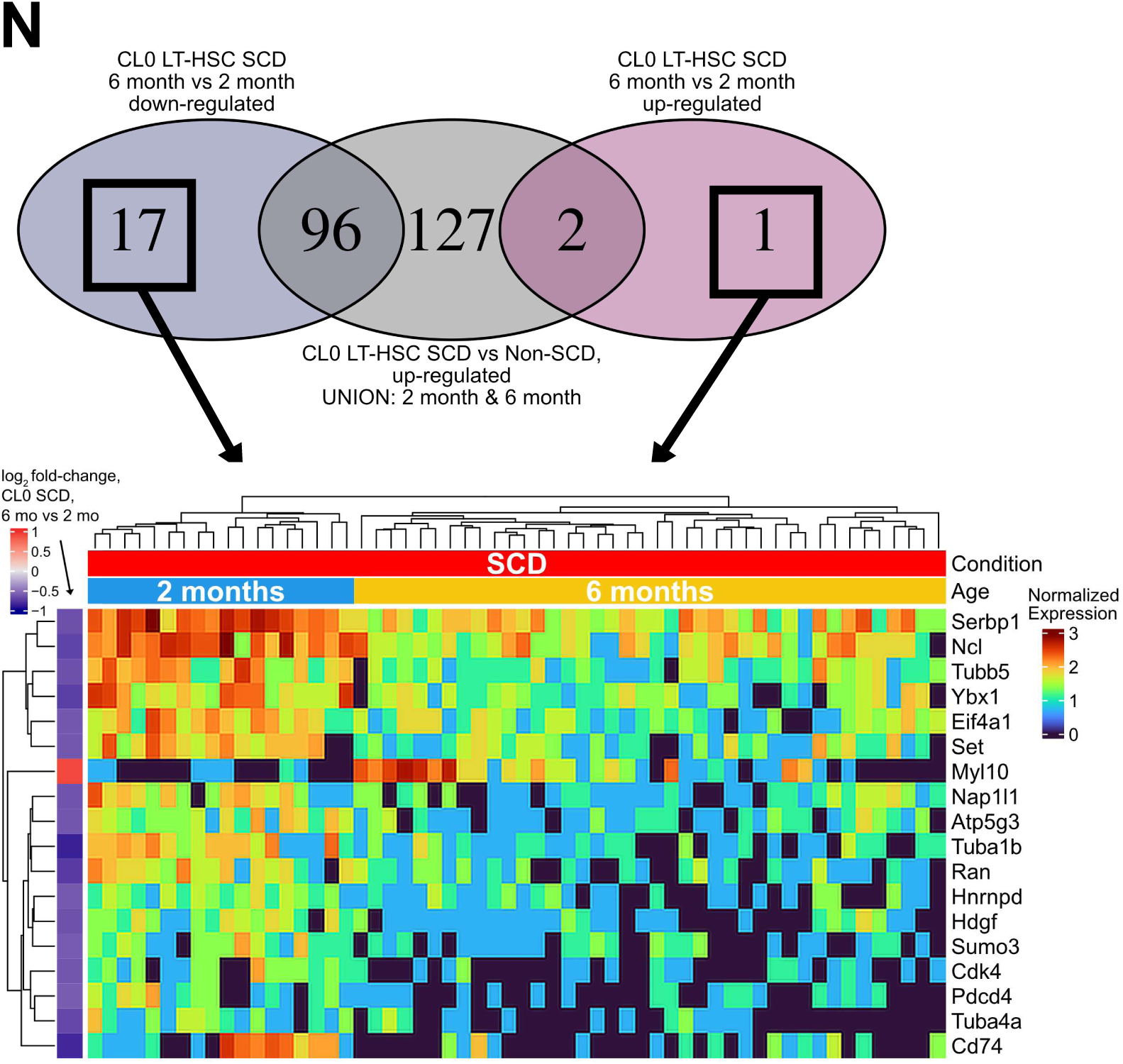
Single cell transcriptional profiling of hematopoietic progenitors isolated from SCD and non-SCD mice - cell type annotation, biological replicate UMAP projection, and differential expression for LT-HSCs. **A**, A tiered ensemble strategy for single-cell level cell type annotation calls for scRNA-Seq. Four well-annotated bone marrow-derived cell type expression datasets were used as references for single-cell level scoring with the R package SingleR. As a secondary approach, cell type marker gene signatures were used for single-cell scoring with AUCell. SingleR and AUCell scores were summarized and combined to identify the highest-scoring cell type per reference for each cell. Considering the Nestorowa et al dataset as a primary reference, cell types from other datasets were assigned when the mean of the alternative references was at least 0.25 higher. All scores were on a [0,1] scale. **B**, Cell types called from the workflow in (A) are concordant with expression of cell type marker genes and cell surface protein markers from CITE-Seq. Each column in the heatmap represents one cell in the single-cell dataset, with the top labels indicating the cell type annotation assigned. SCD or non-SCD and two-month or six-month are indicated by the black/red and yellow/blue color bars, respectively. Antibody-derived tag (ADT) expression for a panel of cell surface protein markers was discretized into “high” and “low” calls by setting a threshold for each ADT separately, indicated by magenta and black, respectively. A set of cell surface marker genes expressed in our dataset were curated from literature and public databases, organized by expected cell type as indicated by the left-side labels. The gene expression heatmap shows z-scores of normalized gene expression across all cells for each gene. **C**, table with total cell counts across the dataset for each annotated cell type, split further by the reference dataset on which the assignment was based (see (A)). The right-side table shows the cell types with fewer than 200 cells in the dataset that were pooled as “other” due to very low representation. Rows of both tables are ordered by the total number of cells per type. “Tab. Muris”: Tabula Muris; “S.Seq”: SMART Seq. **D**, hscScore scores also support the annotated LT-HSCs, in addition to the cell surface markers and marker gene expression. Cell type marker expression from (B) were zoomed in to show the most primitive HSC populations identified in the dataset. Clusters from unsupervised Louvain clustering are also shown. **E**, (i) UMAP projection of c-Kit^+^ bone marrow cells from young and middle-aged SCD and non-SCD mice highlighting distinct clustering of conditions in UMAP space. n=4 biological replicates per condition, with the following pooled cell counts: 2-month-old non-SCD, 14,973 (ii); 2 month-old SCD, 27,149 (iii); 6 month-old non-SCD, 16,182 (iv); 6 month-old SCD, 19,427 (v). In (iv) and (v), black dashed lines indicate clusters composed almost exclusively of cells from SCD animals, and in (iv) pink dashed lines indicate clusters specific to 2 month-old SCD. A cluster was considered “exclusive” to a sample group if other sample groups contributed <5% of cluster cells. **F**, Unsupervised clustering and UMAP projection in two-dimensional space of each biological replicate shown separately. **G**, Fraction of each cell type predicted to be actively cycling (*e.g.* S, G2, or M phase) from Seurat scoring based on cell cycle marker gene expression. Only types with significant differences are shown. Stars indicate FDR-adjusted p-value < 0.05 (omnibus ANOVA followed by post-hoc pairwise t-tests), each point is a biological replicate. **H**, Functional enrichment among pathway-based gene sets for DEGs between LT-HSCs in different clusters, compared to the respective population (e.g. SCD or non-SCD) in cluster 0. The size of each circle indicates the enrichment p-value (for results with FDR-adjusted GOSeq p-values < 0.05) and the color indicates the median log_2_ fold-change for the genes in the intersect, for a given comparison. The top 50 results are shown. **I**, Significantly over-represented pathways (left) and Gene Ontology terms (right) for DEGs from comparison of LT-HSCs between SCD and non-SCD samples within clusters with representation of both conditions at a given age using MAST. The comparisons are indicated at the bottom of each column; rows indicate the pathway or gene set with which the size of the intersect with the DEGs was larger than expected by chance. The size of each circle indicates the enrichment p-value (for results with FDR-adjusted GOSeq p-values < 0.05), and the color indicates the median log_2_ fold-change for the genes in the intersect, for a given comparison. **J**, Left: heatmap of the 92 genes commonly differentially expressed at both 2 and 6 months when comparing SCD vs non-SCD LT-HSCs from cluster 0 (CL0). Right: heatmap of 128 genes differentially expressed in the 2-month comparison but not at 6 months. In both heatmaps, rows and columns are ordered by hierarchical clustering. The left side bars show the log_2_ fold-change for the gene in the indicated comparison. **K**, Ribosomal proteins with elevated gene expression in SCD, which is more pronounced in early progenitors, and not observed in erythroid or myeloid lineage cells or granulocytes. Differential expression of ribosomal protein genes was found through both pseudobulk and single-cell approaches, across multiple cell types, in comparisons between SCD and non-SCD cells. To illustrate the expression patterns of these genes more broadly, expression is shown across all cells for 82 Rps/Rpl genes/pseudogenes with expression in at least 50% of the cells. Rows (genes) and columns (cells) are ordered by hierarchical clustering; clustering of cells was performed within each cell type. **L,** Venn diagram of scDEGs from LT-HSCs of cluster 0 between middle-aged and young samples, separated into up-regulated and down-regulated sets. Only eight DEGs were identified between 6-month and 2-month in non-SCD samples, while 116 were identified between 6-month and 2-month in SCD, with the majority of those having lower expression at 6 months (MAST FDR p-value < 0.05, |fold-change| ≥ 1.5). **M**, Over-represented pathways (left) and Gene Ontology terms (right) for DEGs from comparison of LT-HSCs between SCD and Non-SCD samples and across age within clusters using MAST. The comparisons are indicated at the bottom of each column; rows indicate the pathway or gene set with which the size of the intersect with the DEGs was larger than expected by chance. The size of each circle indicates the enrichment p-value (for results with FDR-adjusted GOSeq p-values < 0.05), and the color indicates the median log_2_ fold-change for the genes in the intersect, for a given comparison. For these comparisons, most genes were up-regulated in SCD vs non-SCD. **N**, 18 genes are modulated only by aging and include important post-translational modifiers and transcriptional regulators, as well as CD74, which can promote HSC maintenance. The left bar indicates the log_2_ fold-change between 6-month and 2-month CL0 SCD LT-HSCs. For pathway-based gene set enrichment in (H), (I), and (M) the source database is indicated by the parentheses, REAC: Reactome, WP: WikiPathways, BioCarta.

**Supplementary Data Figure 7.**
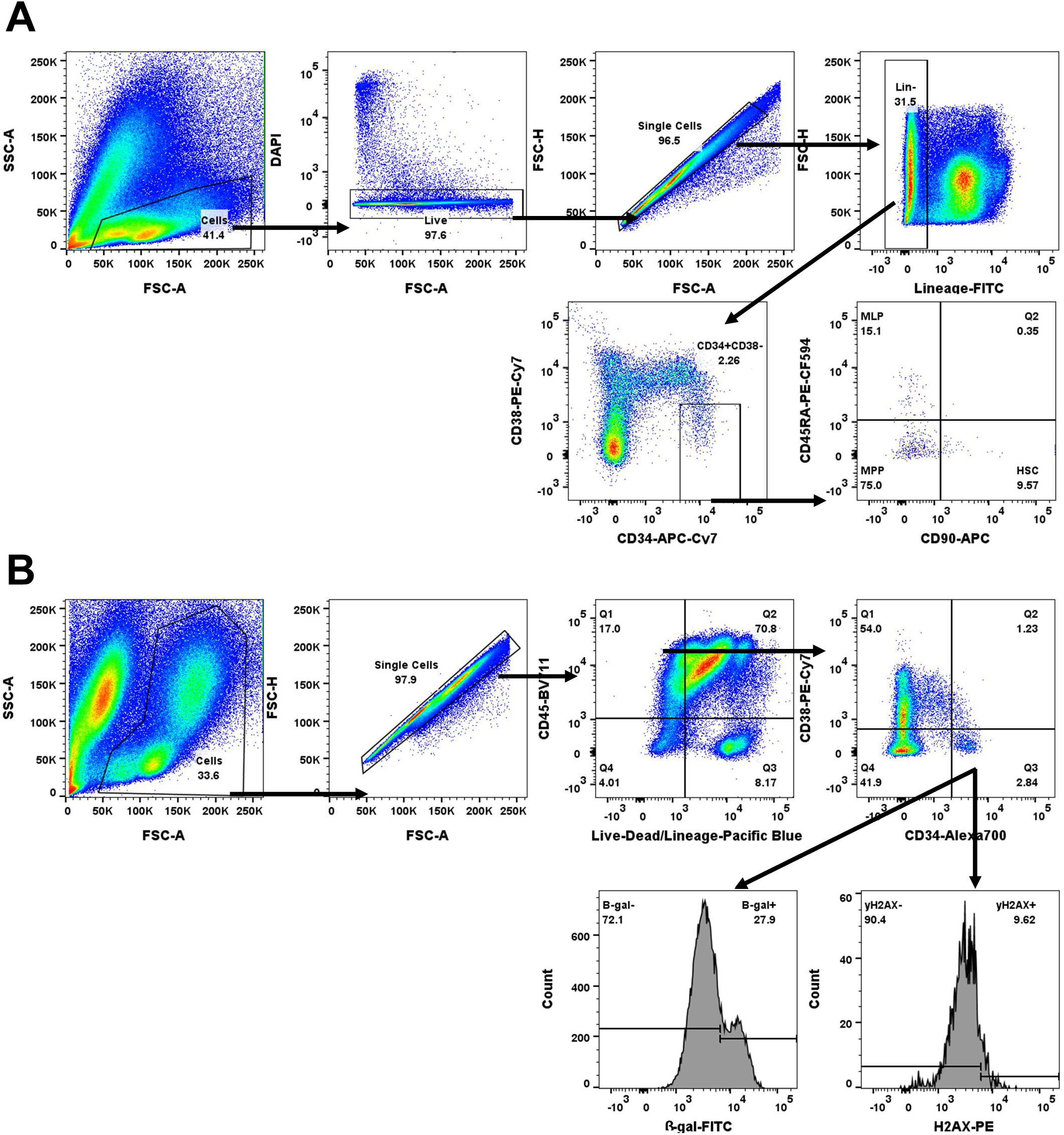
Flow cytometry gating strategies for human bone marrow **A**, Flow cytometry gating strategy for identification of HSCs (Lineage^-^CD34^+^CD38^-^ CD90^+^CD45RA^-^), MPPs (Lineage^-^CD34^+^CD38^-^CD90^-^CD45RA^-^), and MLPs (Lineage^-^CD34^+^CD38^-^CD90^-^CD45RA^+^) in bone marrow of children and young adults with SCD and age-matched non-SCD individuals. **B**, Flow cytometry gating strategy for identification of HSPCs (Lineage^-^CD34^+^CD38^-^) with DNA damage (γH2AX) and intracellular ROS in bone marrow of children and young adults with SCD and age-matched non-SCD individuals.

**Supplementary Data Figure 8.**
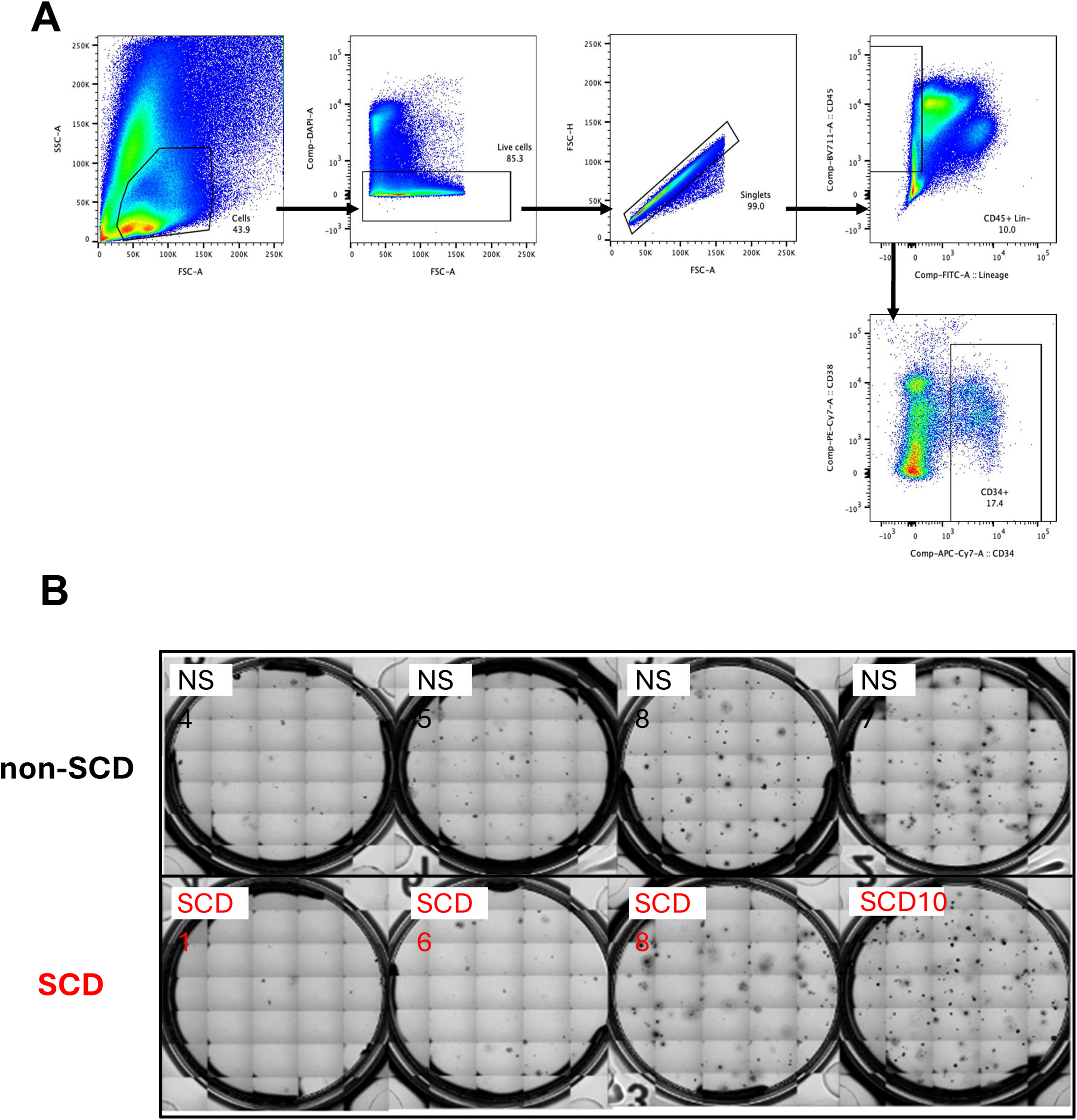
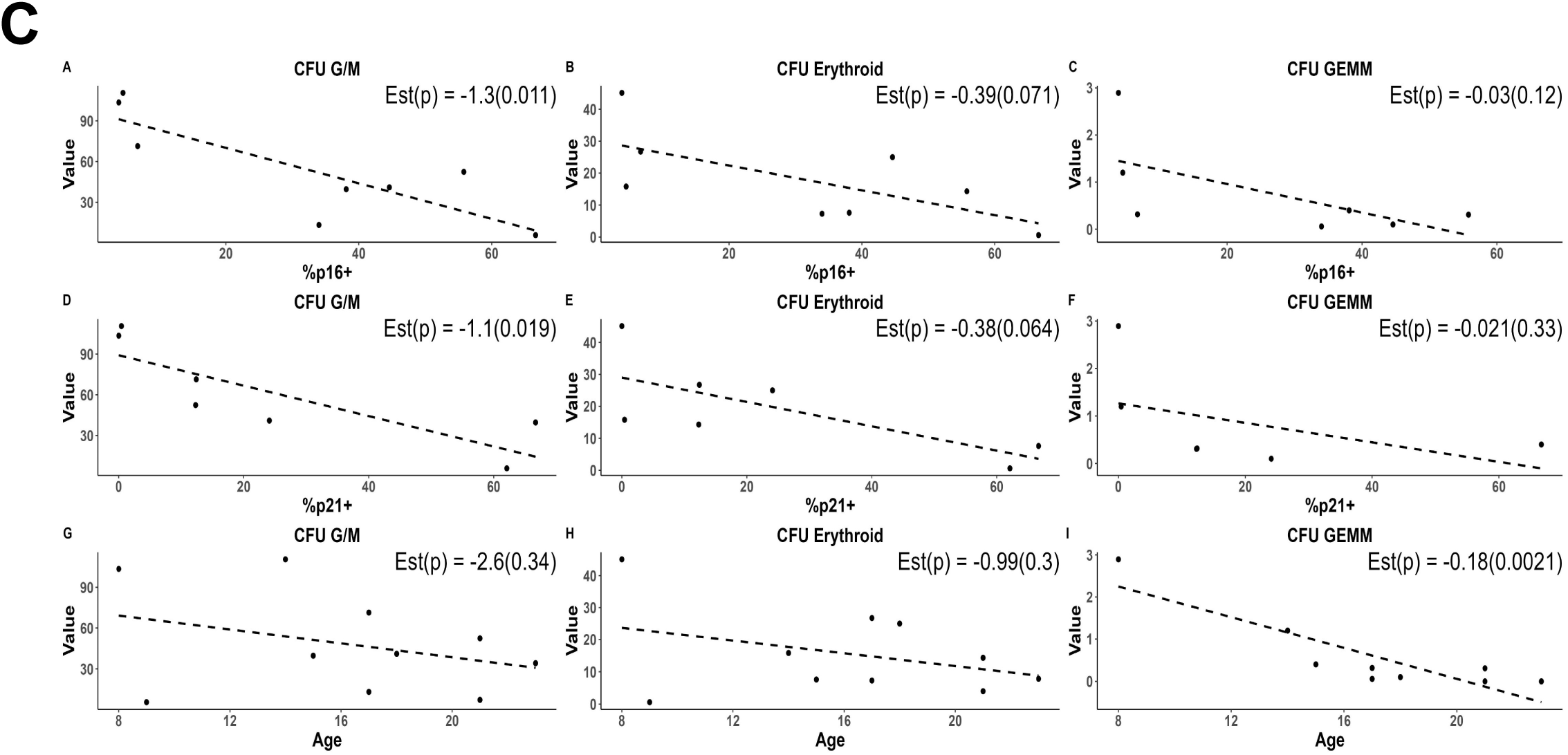
Representative sorting gating strategy and CFU images **A**, Representative FACS gating for isolation of CD45^+^Lineage^-^CD34^+^ for bulk CFU assays. **B**, Representative CFU images from 500 CD45^+^Lineage^-^CD34^+^ cells isolated from non-SCD and SCD individuals. **C**, Correlations of senescence markers and age with total and lineage specific HSPC colony forming ability using a generalized linear mixed model (GLMM) with quasi-Poisson modeling.

**Supplementary Data Figure 9.**
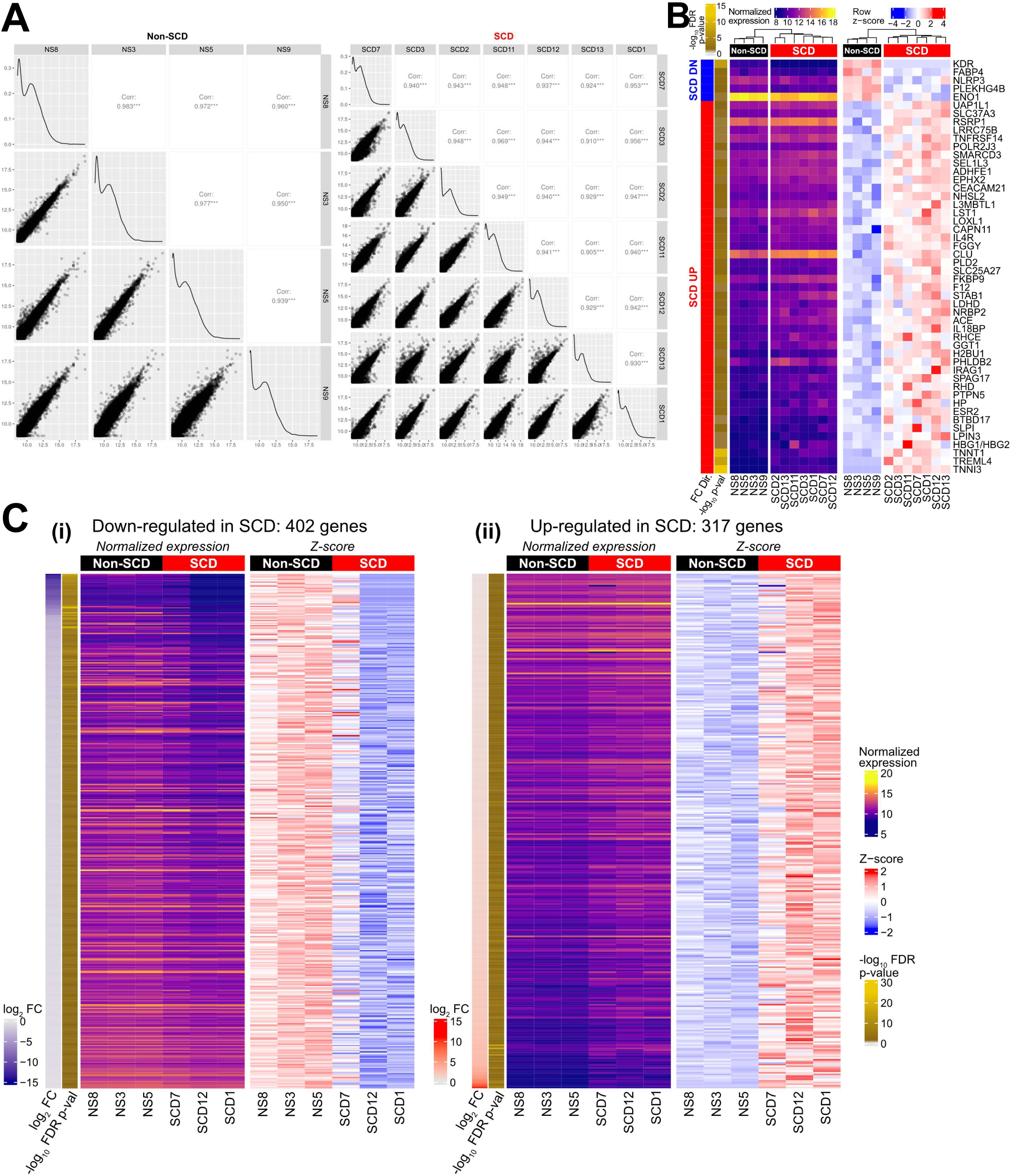
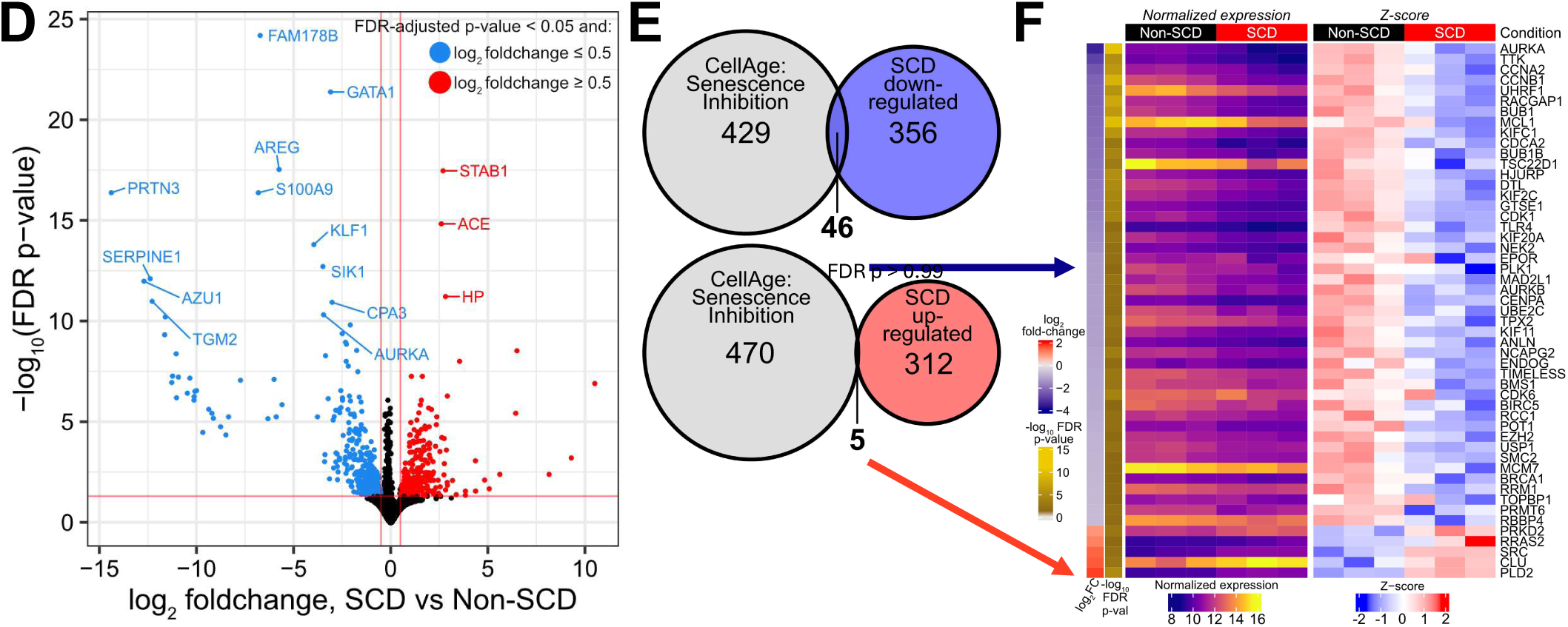
Human Bulk RNA-Sequencing of HSPCs from individuals with and without SCD demonstrates high variability in gene expression between SCD individuals. **A**, Pairwise correlation between biological replicates for individuals without SCD (n = 4, left) and with SCD (n = 7, right) across all expressed coding genes (15,699). The lower triangle shows the scatterplot for each unique pair of samples, the diagonal shows the density of expression for the sample in the indicated column, and the upper triangle shows the Pearson correlation coefficient. Correlation p-values are <0.001 in all cases, indicated by “***”. Pairwise correlations are lower overall between profiles for individuals with SCD than those without SCD. **B**, Heatmap of DEGs between sample groups across all samples using moderate differential expression criteria (FDR-adjusted p-value < 0.1, |fold-change| ≥ 1.5 (0.58 LFC)). Left side: VST-normalized expression for each sample; right side: z-score across rows (genes) to highlight the trends in expression across samples. Samples are grouped by hierarchical clustering, which was performed independently for the expression and z-score heatmaps. The left side-bars indicate the overall direction in fold-change (“FC Dir.”) for the gene (up- or down-regulated in SCD vs non-SCD) and –log10 transformed p-value. Genes are ordered by fold-change. **C**, Heatmap for all DEGs between sample groups for the three most consistent biological replicates within each group, for genes **(i)** down-regulated and **(ii)** up-regulated in individuals with SCD vs non-SCD (FDR-adjusted p-value < 0.05 and log_2_ fold-change < −0.5 or > 0.5, respectively). For each heatmap, genes are ordered by log2 fold-change magnitude, and samples are grouped by hierarchical clustering. For each set of heatmaps, left side: VST-normalized expression for each sample; right side: z-score across rows (genes) to highlight the trends in expression across samples. **D**, Volcano plot from DEG analysis for the samples in (B). Significant DEGs: FDR-adjusted p-value < 0.05, log2 fold-change magnitude ≥ 0.5. 402 genes were down-regulated (blue), 317 were up-regulated (red) in SCD. Red horizontal and vertical lines indicate p-value and fold-change thresholds, respectively. The top 15 significant genes by p-value are labeled. **E**, Gene set enrichment with senescence-associated sets from multiple databases identified significant overlap between SCD-downregulated genes and “senescence inhibition” (GOSeq FDR-adjusted p-value = 0.01) but not SCD-upregulated genes (FDR p-value > 0.99). **F**, Expression of the 51 SCD DEGs intersecting with the CellAge Senescence Inhibition gene set (E). Genes (rows) are ordered by log2 fold-change (left-most side bar), samples are ordered by hierarchical clustering. Left side: VST-normalized expression; right side: row-wise z-scores highlight trends in normalized expression across samples.

---

**Supplementary Data Table 1:** Characteristics of participants included in our analyses.

| Sample ID | Sample Type | Age | Gender | VOC Events* | Acute Chest Syndrome** | Hydroxyurea Duration | Chronic Transfusion*** |
| --- | --- | --- | --- | --- | --- | --- | --- |
| NS1 | Non-SCD | 12 | Male | N/A | N/A | N/A | N/A |
| NS2 | Non-SCD | 21 | Female | N/A | N/A | N/A | N/A |
| NS3 | Non-SCD | 12 | Male | N/A | N/A | N/A | N/A |
| NS4 | Non-SCD | 11 | Female | N/A | N/A | N/A | N/A |
| NS5 | Non-SCD | 9 | Male | N/A | N/A | N/A | N/A |
| NS6 | Non-SCD | 18 | Female | N/A | N/A | N/A | N/A |
| NS7 | Non-SCD | 9 | Female | N/A | N/A | N/A | N/A |
| NS8 | Non-SCD | 15 | Male | N/A | N/A | N/A | N/A |
| NS9 | Non-SCD | 7 | Female | N/A | N/A | N/A | N/A |
| NS10 | Non-SCD | 16 | Male | N/A | N/A | N/A | N/A |
| NS11 | Non-SCD | 2 | Male | N/A | N/A | N/A | N/A |
| NS12 | Non-SCD | 2 | Male | N/A | N/A | N/A | N/A |
| SCD1 | SCD | 9 | Male | 3 | 2 | 8 years 3 months | No |
| SCD2 | SCD | 21 | Male | 3 | 3 | 14 years 10 months | Yes |
| SCD3 | SCD | 17 | Female | 1 | 1 | 14 years 9 months | No |
| SCD4 | SCD | 23 | Male | 9 | 5 | 20 years 5 months | Yes |
| SCD5 | SCD | 15 | Male | 3 | 1 | 12 years 7 months | No |
| SCD6 | SCD | 18 | Male | 2 | 2 | 16 years 3 months | No |
| SCD7 | SCD | 21 | Female | 0 | 0 | 2 months | Yes |
| SCD8 | SCD | 17 | Female | Unknown | 2 | 13 years 6 months | Yes |
| SCD9 | SCD | 14 | Male | 0 | 2 | 7 years 11 months | No |
| SCD10 | SCD | 8 | Male | 0 | 0 | 6 years | Yes |
| SCD11 | SCD | 13 | Male | 9 | 5 | 7 years 11 months | No |
| SCD12 | SCD | 17 | Male | 7 | 1 | 15 years 3 months | No |
| SCD13 | SCD | 16 | Male | 2 | 4 | 3 years 6 months | No |
| SCD14 | SCD | 6 | Female | 4 | 1 | 5 years 6 months | No |
| SCD15 | SCD | 6 | Male | 0 | 2 | 5 years 8 months | No |
| SCD16 | SCD | 15 | Female | 4 | 0 | 6 years 3 months | No |

\*Vaso-occlusive (VOC) events refer to any acute pain crises experienced by the participants in the 2 years prior to enrollment in the clinical study.
\*\*Acute chest syndrome refers to any acute chest syndrome events experienced by the participants in the 2 years prior to enrollment in the clinical study.
\*\*\*Chronic transfusion refers to the regular receipt of blood transfusions approximately every 3-5 weeks to treat symptoms of severe SCD.

